# FMRP regulates mRNAs encoding distinct functions in the cell body and dendrites of CA1 pyramidal neurons

**DOI:** 10.1101/2021.07.18.452839

**Authors:** Caryn R. Hale, Kirsty Sawicka, Kevin Mora, John Fak, Jin Joo Kang, Paula Cutrim, Katarzyna Cialowicz, Thomas Carroll, Robert B. Darnell

## Abstract

Neurons are believed to rely on dendritic localization and translation of mRNAs in order to generate activity-dependent changes in the synaptic plasticity. Here, we develop a strategy combining compartment-specific CLIP and TRAP in conditionally tagged mice to precisely define the ribosome-bound dendritic transcriptome of CA1 pyramidal neurons. This revealed transcripts that have differentially localized alternative 3’UTR and splicing isoforms. FMRP targets are overrepresented among dendritic mRNAs, and compartment-specific FMRP-CLIP defined 383 dendritic FMRP targets, and also allowed for segregation of whole-cell FMRP targets into functional modules that are locally regulated by FMRP. In the absence of FMRP, dendritic FMRP targets show increased ribosome association, consistent with reported roles for FMRP in translational repression. Together, the data support a model in which distinct patterns of FMRP localization allow it to differentially regulate the expression of nuclear proteins and synaptic proteins within different compartments of a single neuronal cell type.

## Introduction

A key feature in the molecular biology of learning and memory is protein-synthesis dependent synaptic plasticity, which involves translation of localized mRNAs in response to synaptic activity. Local translation has been demonstrated in neuronal dendrites (reviewed in (Glock et al., 2017)) and axons (reviewed in (Lin and Holt, 2007; Rangaraju et al., 2017)) and allows for rapid and precise changes in the local proteome near active synapses. In dendrites, a brief burst of local translation has been shown to be necessary and sufficient for induction of the late phase of long-term potentiation (L-LTP, occurring hours to days after potentiation) (Frey et al., 1988; Kang and Schuman, 1996; Kang et al., 1997) and long-term depression (LTD) (Huber, 2000), and inhibiting protein synthesis blocks long-term memory formation (Sutton and Schuman, 2006).

Activity-dependent local translation depends on both the availability of specific mRNAs and the sensitivity with which their translation can be initiated upon local signaling events. Both rely on interactions between mRNAs, a host of RNA-binding proteins, and ribosomes. mRNAs are thought to be transported in a translationally repressed state into the neuronal processes via transport granules containing RNA-binding proteins such as FMRP, CPEB1, ZBP-1 and Stau1/2 (Hüttelmaier et al., 2005; Krichevsky and Kosik, 2001; Martin and Ephrussi, 2009). Although dendritic targeting elements have been defined for a few mRNAs such as *CamkIIα*, *β-actin*, and *Map2* (Andreassi and Riccio, 2009), the functional relationship between the dendritic transcriptome and these RNA-binding proteins is still largely unknown. For at least some localized mRNA granules, signaling cascades initiated by synaptic activity lead to their dissolution and initiation of translation (Dahm and Kiebler, 2005), but the role of RNA regulatory factors in this process is incompletely understood. The integrated study of the dendritic transcriptome and the RNA-binding proteins responsible for mRNA localization and regulation of local translation will provide critical insight into mechanisms underlying protein-synthesis dependent synaptic plasticity.

Fragile X mental retardation protein (FMRP), the RNA-binding protein whose activity is lost in Fragile X Syndrome, represses translation (Bassell and Warren, 2008; Costa-Mattioli et al., 2009; Darnell et al., 2011; Laggerbauer et al., 2001) and is thought to be a key regulator of activity-dependent local translation in neurons (Banerjee et al., 2018; Bear et al., 2004; Huber et al., 2002; Lee et al., 2011). Dendritic FMRP levels are increased upon neuronal activity, with evidence for local translation of the FMRP transcript itself (Greenough et al., 2001; Weiler et al., 1997) and kinesin-mediated movement of FMRP-containing mRNA transport granules from the neuronal cell body (Dictenberg et al., 2008). At the synapse, FMRP is proposed to be linked to local signal transduction, potentially through calcium-regulated post-translational modification of the protein, which alters the FMRP granule and leads to translation of the mRNAs (Lee et al., 2011; Narayanan et al., 2007). FMRP knockout (KO) neurons show excess basal translation as well as an inability to produce activity-stimulated translation (Ifrim et al., 2015).

FMRP targets have been described in the whole mouse brain (Darnell et al., 2011; Korb et al., 2017) and, using TRAP (Ceolin et al., 2017; Kumari and Gazy, 2019), or through TRAP together with CLIP, (Sawicka et al., 2019), specifically in the excitatory CA1 neurons of the mouse hippocampus. FMRP target genes in these neurons overlap significantly with autism susceptibility genes and include genes involved in both synaptic function and transcriptional control in the nucleus (Darnell, 2020; Darnell et al., 2011; Iossifov et al., 2012; Sawicka et al., 2019), and loss of FMRP increases translation of chromatin modifiers such as BRD4 (Korb et al., 2017) and SETD2 (Shah et al., 2020). These and other observations have suggested a model in which FMRP regulates the stoichiometry of its targets in two ways: globally, by translational control of transcription regulators in the cell body, and locally, by enabling activity-dependent local translation of synaptic proteins in dendrites (Darnell, 2020), but it is still unclear how these occur simultaneously in a single neuron. Here, we probe this model by exploring the subcellular localization of FMRP and specific FMRP-bound mRNAs in CA1 neurons of wild-type and *Fmr1*-null mice, examining whether together they may dictate RNA localization and translation.

We utilize compartment- and cell-type specific profiling technologies to precisely define the dendritic transcriptome of mouse hippocampal CA1 pyramidal neurons. RNA profiling of these compartments reveals that dendritic mRNAs are enriched for elongated 3’UTR isoforms and depleted for alternative splicing (AS) events driven by the neuronal splicing factor NOVA2, indicating a nuclear role in the generation of the localized transcriptome in CA1 neurons. Integrating compartment-specific cTag-FMRP-CLIP and TRAP defined FMRP CLIP scores in the dendrites and cell bodies of CA1 neurons and identified 383 FMRP-bound dendritic targets. This allowed us to distinguish FMRP targets according to their site of regulation within neurons, revealing enrichment of FMRP-regulated mRNAs encoding nuclear proteins in the CA1 cell bodies, and mRNAs encoding synaptic proteins in the CA1 dendrites. Moreover, mRNAs encoding these synaptic proteins show altered ribosome association in FMRP KO mice. Together these findings support a model in which distinct patterns of both mRNA and FMRP subcellular localization allow FMRP to regulate the expression of different proteins within different compartments in a single neuronal cell type.

## Results

### Identification and characterization of the dendritic transcriptome in hippocampal pyramidal neurons *in vivo*

We developed a system that allows for parallel isolation of mRNAs and RNA-binding proteins that are enriched in the cell bodies or dendrites specifically in excitatory CA1 neurons in the hippocampus (Figure 1A). We created three cell-type specific conditionally protein-tagged mouse lines by crossing either the RiboTag (Sanz et al., 2009), cTag-PABP (Hwang et al., 2017), or Fmr1-cTag (Sawicka et al., 2019; Van Driesche et al.) mouse lines with mice in which Cre recombinase expression is driven from the CamkIIα promoter (Tsien, 1998). This results in mouse lines expressing tagged ribosomes, or in the case of polyA-binding protein c1 (PABPC1) or FMRP, “knock-in” tagged proteins expressed from native genes, specifically in the *CamkIIα*-expressing CA1 pyramidal neurons of the mouse hippocampus (Figure S1A). Therefore, microdissection of the CA1 neuropil compartments enriches for dendritic tagged proteins that originated from the cell bodies (CB) of the CA1 neurons. Affinity purification of tagged proteins enriches for mRNAs bound to localized ribosomes (in CamKIIa-Cre x RiboTag mice by TRAP), PABPC1 (in CamKIIa-Cre x cTag-PABP by PAPERCLIP) or FMRP (in CamKIIa-Cre x Fmr1-cTag mice by FMRP-CLIP) and de-enriches for mRNAs from the glia or interneurons that are also present in the neuropil.

**Figure 1.**
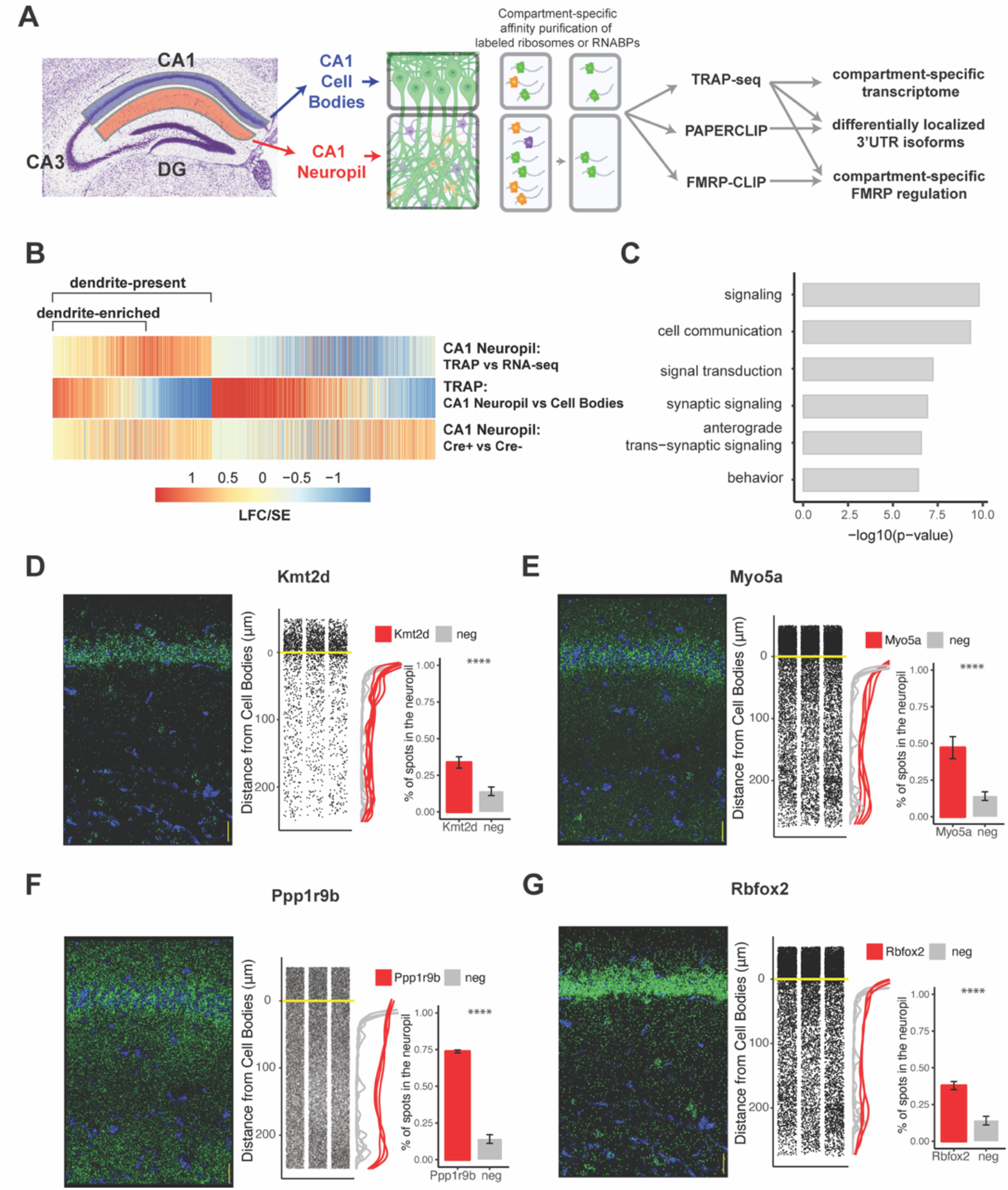
Combining cell-type specific protein tagging and manual microdissection allows for precise definition of the CA1 dendritic transcriptome. A) *Experimental Design.* Hippocampal slices from CamkIIa-Cre expressing conditionally tagged mouse lines were subject to microdissection in order to separate the Cell Bodies and CA1 neuropil layers. These layers contained material from pyramidal neurons (in which proteins of interest contain an affinity tag) and contaminating cell types. Microdissected compartments were subject to affinity purification in order to obtain pyramidal neuron-specific ribosomes or affinity-tagged RNA-binding proteins and bound mRNAs. In order to obtain the dendritic ribosome-bound transcriptome, TRAP-seq was performed from tagged ribosomes in the CA1 neuropil compartment. Compartment-specific cTag-PAPERCLIP was performed in order to determine mRNAs with 3’UTR isoforms that undergo differential localization, and compartment-specific FMRP regulation was determined by cTag FMRP-CLIP of the microdissected compartments. B) *Identification of dendritic mRNAs.* Differential gene expression was performed on bulk RNA-seq and TRAP-seq from microdissected CA1 compartments. Colors indicate the log2 fold change (LFC)/SE (standard error, stat) from DESeq analysis. mRNAs significantly enriched in CA1 neuropil-TRAP over bulk RNA-seq of the CA1 neuropil were defined as “dendrite-present”. mRNAs that were also significantly enriched in CA1 neuropil TRAP when compared to Cell Bodies TRAP were considered to be “dendrite-enriched”. In addition, only mRNAs that were enriched in CA1 neuropil TRAP in Camk-Cre expressing RiboTag mice when compared to RiboTag mice not expressing Cre were considered. C) *Localized mRNAs are highly enriched for genes involved in synaptic signaling and synapse organization.* GO analysis was performed comparing dendrite-enriched mRNAs to all mRNAs expressed in CA1 neurons. D-G) *Validation of localized mRNAs.* FISH was performed using the RNAscope method using probes designed against the entire mRNA of the indicated gene. *(left)* Representative FISH image of RNAscope on the CA1 region of coronal brain sections. mRNA spots are shown in green, and DAPI staining is shown in blue. Scale bars represent 30 microns. (middle) The distance between the mRNA punctae and the Cell Bodies were quantitated for three representative images. Density is plotted for all collected images (red) and compared to an mRNA (Snca) that was identified as sequestered in the Cell Bodies (grey). (right) mRNAs more than 10 microns from the Cell Bodies were considered to be in the neuropil. The percent of indicated mRNAs that were found in the neuropil is plotted. A Cell Body-sequestered mRNA (Snca) is used as a negative control. Stars indicate results of the Wilcoxon ranked test. (**** indicates p < .00001).

Microdissected CA1 compartments from 8-10 week old mice were subjected to bulk RNA-seq as a denominator for all transcripts in the neuropil, and TRAP as a denominator for all CA1 pyramidal neuron-specific, ribosome-bound dendritic transcripts. Immunoprecipitation (IP) conditions were optimized to isolate relatively pure, intact, ribosome-bound mRNAs with minimal contamination by interneurons and glial cells found in the neuropil (Figure S1B-E). As a negative control, animals not expressing the Cre recombinase were microdissected and subject to affinity purification and sequencing, and only mRNAs enriched over controls were considered for downstream analyses. We identified two groups of dendritic mRNAs: dendrite-present (significantly enriched in CA1 neuropil TRAP-seq over CA1 neuropil bulk RNA-seq; 2058 mRNAs) and dendrite-enriched (dendrite-present and significantly enriched in CA1 neuropil TRAP over cell bodies TRAP; 1211 mRNAs; Figure 1B, see Supplemental Tables 1-2 for a full list of mRNAs identified). 689 (34%) of the dendrite-present mRNAs were previously identified in bulk RNA-seq of the microdissected rat CA1 neuropil (Cajigas et al., 2012), of which 334 (48%) are also in the dendrite-enriched group (Figure S2A-B).

Our identified dendrite-enriched mRNAs are significantly longer than the whole-cell transcriptome for pyramidal neurons in the CA1 (Sawicka et al., 2019), whether considering full length transcripts, 5’UTR, 3’UTR or coding sequence (CDS) portions (Figure S2C). Gene Ontology (GO) analysis of dendrite-enriched mRNAs shows strong enrichment for genes encoding proteins with important roles in the synapse such as synaptic signaling, anterograde synaptic signaling and behavior (Figure 1C), consistent with prior analyses (Cajigas et al., 2012). We validated the localization of several mRNAs using RNAscope Fluorescence In situ Hybridization (FISH), that had not been identified in previous studies including *Kmt2d, Myo5a*, *Ppp1r9b,* and *Rbfox2* (Figures 1D-1G). Interestingly, approximating the distance traveled from the cell body for each detected mRNA spot reveals variable mRNA distribution patterns for different transcripts, suggesting multiple potential paths for mRNA localization. For example, roughly 35% of the transcripts encoding *Kmt2d* and *Rbfox2* were detected throughout in the neuropil, whereas ∼74% of the transcripts encoding *Ppp1r9b* were highly localized to the distal neuropil (Figure 1F). By comparison, less than 15% of *Snca* mRNAs, which we identified as enriched in the CA1 cell body compartment, were found in the CA1 neuropil (data not shown).

### Identification of mRNAs with 3’UTR isoforms preferentially localized to dendrites

Subcellular localization of cytoplasmic mRNAs is thought to be at least partially mediated by 3’UTR elements (Andreassi and Riccio, 2009; Blichenberg et al., 2001; Mayford et al., 1996; Tushev et al., 2018). However, analysis of 3’UTRs from RNA-seq data alone is complicated by mixed cell types, incomplete annotation and difficulty in identifying internal polyA sites. To identify the expressed 3’UTRs in CA1 pyramidal neurons, we combined polyA sites determined by CamkIIα-Cre driven cTag-PAPERCLIP from whole hippocampus (Hwang et al., 2017) and microdissected CA1 compartments (Figure S3) with splice junctions detected in pyramidal neuron-specific TRAP, in order to identify the boundaries of potential 3’UTRs (Figures 2A and S4A-B). Combining these datasets revealed 15,322 3’UTRs expressed in CamkIIα-expressing pyramidal neurons, including 3,700 genes that give rise to mRNAs with more than one 3’UTR isoform. Analyzing expression of these 3’UTRs in the compartment-specific TRAP data revealed 219 3’UTR isoforms that were differentially localized to CA1 dendrites (Figure 2B, Supplemental Table 4).

**Figure 2.**
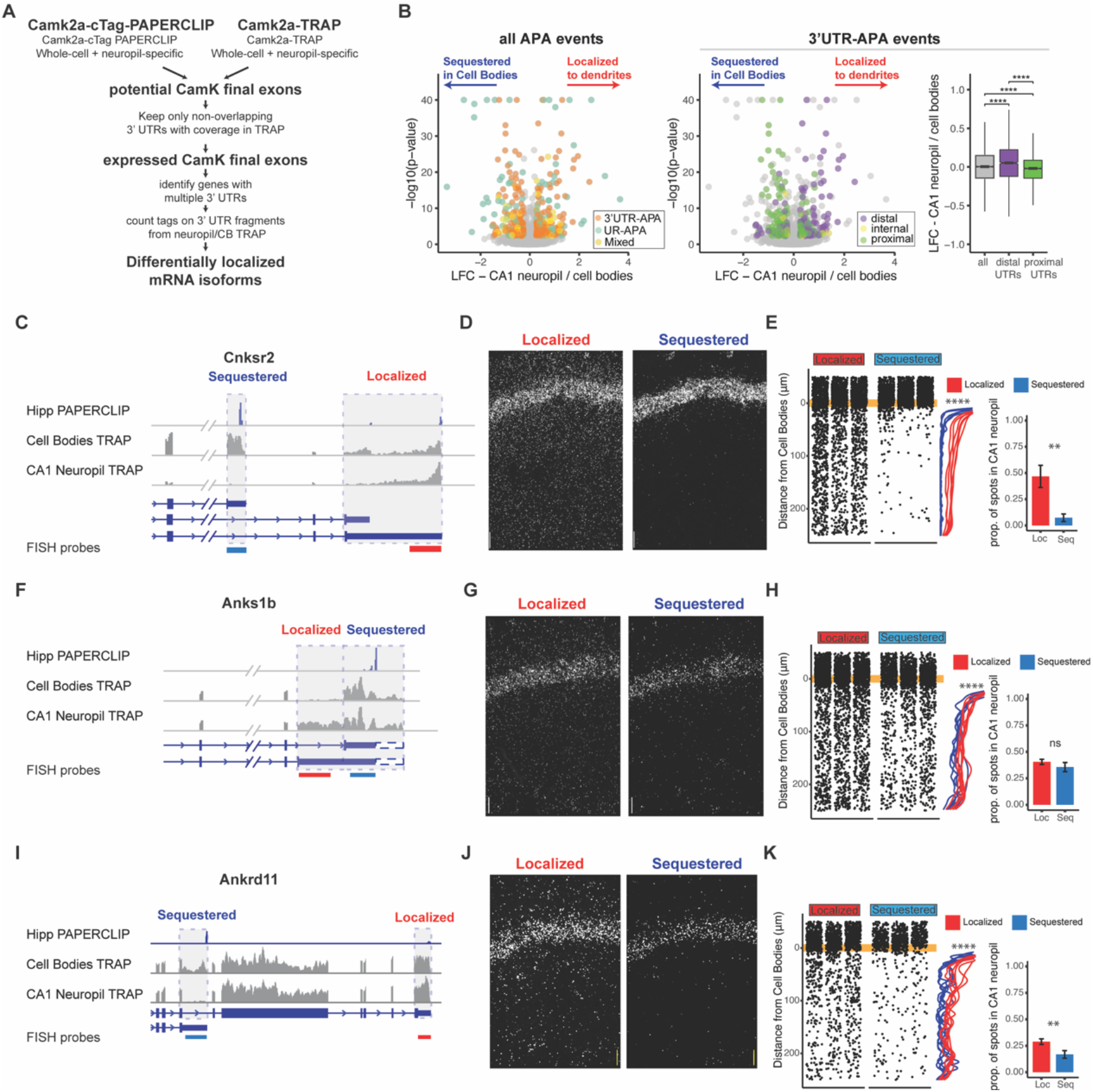
Combining cTag PAPERCLIP and TRAP in order to identify genes with differentially localized 3’UTR isoforms. A) *Scheme for identification of expressed 3’UTR isoforms in CA1 neurons followed by analysis of differential localization*. Boundaries of expressed 3’UTR isoforms in CA1 neurons were determined by combining polyA sites, determined by cTag-PAPERCLIP from both the whole hippocampus and microdissected CA1 compartments, with splice junctions from cell-type specific TRAP experiments. These potential final exons were filtered for non-overlapping 3’UTRs with complete coverage in TRAP. Compartment-specific expression of the resulting 3’UTRs was quantitated, and DEX-seq was used to determine 3’UTRs that were differentially localized in the dendrites of CA1 neurons. B) *Differential localization of 3’UTR isoforms.* (left) Volcano plot shows LFC (calculated by DEXSeq) vs. FDR for genes that produce significant differentially localized 3’UTR isoforms, colored by types of APA events. 3’UTR-APA (orange) are 3’UTRs with multiple polyA sites, and APA changes do not affect the CDS of the resulting mRNAs. Genes that undergo Upstream region APA, or UR-APA (green), utilize polyA sites within introns upstream of the 3’UTR, and result in mRNAs with truncated CDS. Genes that undergo both types of APA are shown in yellow. (middle) 3’UTR-APA events are colored according to their position in the gene, either proximal to the stop codon (green), internal (yellow), or distal (purple). (right) For all 3’UTR-APA events detected, distal 3’UTRs (purple) are significantly enriched in the CA1 neuropil (determined by the LFC of CA1 neuropil / Cell Bodies), when compared to proximal (green) 3’UTRs. p-values for paired Wilcoxon ranked tests are indicated (**** = p < .00001). C-E) *Validation of differential localization of 3’UTR isoforms of Cnksr2 isoforms*. C) Distribution of Cell Bodies- and CA1 neuropil TRAP-seq reads for the 3’ end of the Cnksr2 mRNA. Camk2a-cTag PAPERCLIP tags from the hippocampus are shown in blue. Coverage is normalized for read depth and scaled in order to best illustrate isoform expression. Predicted mRNAs are indicated below, and the positions of the FISH probes are indicated (the sequestered probe is shown in blue, and the localized probe in red). D) smFISH on the CA1 region using probes against localized (left) and sequestered (right) 3’UTR sequences. E) (left) Spots were counted in either the CB or CA1 neuropil region and distance traveled from the cell body was determined for each spot. Plots show location of spots in all quantitated replicates. Line plots show the density of the detected spots that were found in the CA1 neuropil in either the Cell Body-sequestered (blue) or neuropil-localized 3’UTR isoform (red) for the 300 nts proximal to the Cell Bodies. Stars indicate significance in kolomogorov-smirnov tests (**** = p-value < .00001) between distribution of sequestered and localized 3’UTR isoforms. (right) Overall quantitation of spots in the Cell Bodies (< 10 μm from the Cell Body Layer) and CA1 neuropil (> 10 μm from the Cell Bodies) is shown in barplots. Results of Wilcoxon ranked sum tests are shown (** = p-value < .001). F-H) *Differential localization of Anks1b 3’UTR isoforms.* See description for C-E. Dashed box indicates a potential underutilized 3’UTR extension that is observed by TRAP, but represents only a minor fraction of PAPERCLIP reads. I-K) *Differential localization of Ankrd11 3’UTR isoforms*.

Analysis of these differentially localized 3’ UTRs revealed transcripts generated by two types of alternative polyadenylation (APA), distinguished by their effect on the coding sequence of the resulting protein. APA events that do not affect the coding sequence of the resulting protein derive from transcripts with multiple polyadenylation sites in a single 3’UTR, resulting in isoforms with short (proximal) and long (distal) 3’UTRs (3’UTR-APA). APA events that truncate the coding sequence of the resulting protein utilize polyA sites in upstream regions, resulting in multiple protein isoforms (UR-APA) (Tian and Manley, 2017). Of the 219 genes producing differentially localized 3’UTR isoforms in CA1 neurons, we found 149 had no effect on the CDS, 48 resulted in altered CDS, and 22 generated both event types (Figure 2B, left panel). Among isoforms with unchanged CDS, distal 3’UTRs were significantly enriched in dendrites, consistent with a previous study of CA1 neuropil RNAs analyzed by 3’ end sequencing (Tushev et al., 2018). Conversely, proximal 3’UTRs were significantly enriched in the CA1 cell bodies (Figure 2B, middle and right panel). We used FISH to validate these types of differential localization events, including Calmodulin 1 (*Calm1*) (Figures S4B-S4E), previously described to harbor differentially localized 3’UTR isoforms (Tushev et al., 2018), F-box protein 31 (*Fbxo31*) (Figures S4F-S4H), an E3 ubiquitin ligase proposed to be involved in neuronal maintenance and dendritic outgrowth (Vadhvani et al., 2013), and vesicle-associated membrane protein B (*Vapb*) (Figures S4I-S4K), a membrane protein involved in vesicle trafficking.

Approximately 20% of the differential isoform localization events (48 out of 219) involved a polyadenylation event that led to an extension or truncation of the coding sequence. For example, the gene for connector enhancer of kinase suppressor of Ras2, (*Cnksr2* or MAGUIN) produces mRNAs with two 3’UTRs isoforms: a short isoform that is highly sequestered in the cell bodies (less than 10% of transcripts were found in the CA1 neuropil by FISH), and a longer isoform of which at least 40% were localized in the CA1 neuropil (Figure 2C-2E). Analysis of the ankyrin repeat and sterile alpha motif domain containing 1B (*Anks1b*) gene reveals differential localization of an isoform generated from 5’ extension of the 3’UTR sequence, which was depleted in the CA1 cell bodies, and again validated by FISH (Figures 2F-2H). Finally, two mRNAs produced from the ank-repeat domain containing protein 11 (*Ankrd11*) gene were identified, a full-length version that contains Ank repeats as well as the C-terminal transcriptional repression and activation domain, and a previously-uncharacterized isoform that results from use of a polyadenylation site found in intron 8 in order to produce a protein that contains only the Ank-repeat regions (see PAPERCLIP profile in Figure 2I). The truncated isoform was predominantly detected in the cell bodies of CA1 neurons by both TRAP and FISH, while the full-length isoform was detected in both the cell bodies and dendrites (Figures 2I-2K). Together, these data demonstrate the utility of combining compartment- and cell-type transcriptomics and PAPERCLIP in order to define expressed 3’UTRs and reveal localized transcripts generated by alternative processing of 3’ UTRs.

### Identification of mRNAs with alternative splicing isoforms that are preferentially localized to the dendrites

We next sought to identify alternative spliced RNA isoforms that were differentially localized to the dendrites of CA1 pyramidal neurons. After identification by rMATS (Shen et al., 2014) and filtering, we identified 165 alternative splicing events in 143 genes that were differentially localized (Figure 3A, Supplemental Table 5). Of these, 106 (64.2%) are skipped exons, 32 (19.4%) are alternative 3’ splice sites, 14 (8.5%) are alternative 5’ splice sites, and 13 (7.9%) are mutually exclusive exons (Figure 3B). These alternatively spliced transcripts encoded proteins involved in synaptic functions such as action potential, receptor localization and synaptic signaling, and in mRNA splicing (Figure 3C).

**Figure 3.**
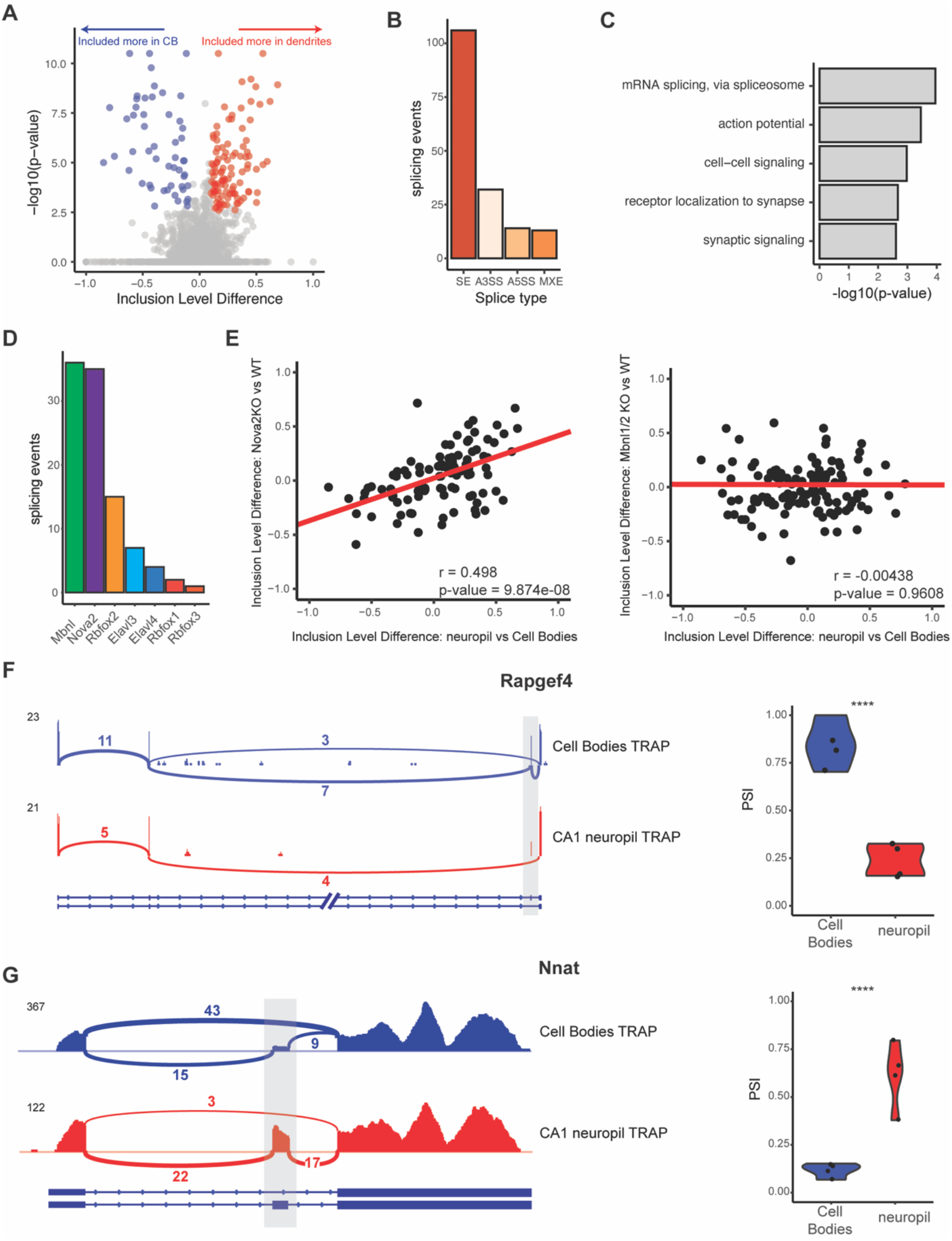
Differential localization of mRNAs with alternative splice events. A) *Analysis of Cell Bodies and CA1 neuropil TRAP by rMATs reveals differentially localized alternative splice events*. Volcano plot shows the Inclusion Level Difference vs. the -log10(p-value) for each detected splice event. Significant events (FDR < .05, |dPSI| > .1) are colored either red (included more in the CA1 dendrites) or blue (included more in Cell Bodies). B) *Types of splicing events identified as differentially localized.* C) *GO analysis reveals enriched functional terms for mRNAs with differentially localized alternative splice events.* All mRNAs expressed in CA1 neurons were used as a background. D) *Neuronal RNA-binding proteins that are responsible for differentially localized AS events*. Alternative splicing analysis was performed on RNA-binding protein KO vs WT RNA-seq data (see Supplemental Table 4 for sources of data). Splicing events that were shown to be differentially localized (seen in A) and also changed in the absence of the RNA-binding protein are plotted. E) *Nova2 neuronal splicing factor generates Cell Body-restricted mRNA transcripts*. Inclusion level differences in CA1 neuropil vs. Cell Bodies-TRAP-seq are compared with the Inclusion level differences in Nova2-/- vs. WT RNA-seq data (left) and Mbnl1/2 -/- RNA-seq (right). Red line indicates a fitted linear model of the data. Results of the Pearson correlation test are shown. F) *Differential localization of spliced Rapgef4 mRNAs*. Representative Sashimi plots (left) are shown for Cell Bodies- (blue) and CA1 neuropil (red) TRAP-seq. Numbers of detected splice junctions are shown. Violin plots (right) show the PSI values for the alternative splice event shown in the Sashimi plot. Each dot represents a single TRAP-seq replicate. Stars indicate significance (FDR) of the splicing change, as determined by rMATs (**** = FDR < .00001). E) *Differential localization of spliced Nnat mRNAs*. (see D)

To determine splicing factors that may be responsible for these differentially localized AS events, we used existing datasets (Supplemental Table 6) of splicing changes previously found to be mediated by neuronal alternative splicing factors. Of these, MBNL1/2 and NOVA2 were found to regulate the largest number of these events (37 for MBNL1/2 and 36 for NOVA2, Figure 3D). Interestingly, we found that CA1 neuropil/cell body splicing changes were positively correlated with splicing changes in NOVA2 KO animals (from analysis of *Nova2*-null vs WT data, pearson coefficient = 0.498, p-value = 9.87e-08), which indicates that NOVA2 drives splicing changes that result in mRNAs that are preferentially sequestered in CA1 cell bodies (Figure 3E, left panel). This effect was specific for NOVA2, as MBNL1/2-dependent splicing changes did not show such a correlation with the localized splicing changes (Pearson coefficient = -0.00438, p-value = 0.9698, Figure 3E, right panel).

Among transcripts that exemplify differential exon usage in localized transcripts were *Rapgef4/Epac2* and neuronatin (*Nnat*). *Epac2*, the gene encoding a cAMP-activated guanine exchange factor for RAP1and RAP2 involved in LTP in the hippocampus, expresses two isoforms (*Epac2A1* and 2A2) that are expressed in the brain (Hoivik et al., 2013). Of the *Epac2A* isoforms detected in the CA1 dendrites, only 25% were the *Epac2A2* isoform, whereas in the cell bodies, 75% of the *Epac2A* transcripts were the *Epac2A1* isoform which does not contain exon 7 (Figure 3F), indicating preferential localization of the *Epac2A2* to the CA1 dendrites. *Nnat*, a maternally-imprinted gene whose protein is important for regulation of intracellular calcium levels, is expressed as either an α- and β-isoform in which exon 2 is included or skipped, respectively. We found that *Nnat-β* is predominantly sequestered in the cell bodies, with only ∼12.5% of cell body transcripts containing exon 2. Conversely, the majority of localized *Nnat* transcripts (50-75%) contain exon 2, indicating preferential localization of the *Nnat-α* subunit (Figure 3G). In sum, these data underscore the role that regulation of alternative splicing can play in localization of specific transcript isoforms.

### CA1 FMRP targets are over-represented in the dendritic transcriptome

FMRP is thought to be a master regulator of local translation (Ronesi and Huber, 2008), leading us to examine the relationship between the dendritic ribosome-bound transcriptome and FMRP binding. We observed significant over-representation of CA1 whole-cell FMRP targets in the dendrite-present and even more so in dendrite-enriched mRNAs (Fig 4A). Of 1211 dendrite-enriched mRNAs, about 35% (413 mRNAs) were FMRP targets, compared to 28.5% of dendrite-present mRNAs and 11.6% of all CA1-expressed mRNAs (Figure 4B).

**Figure 4.**
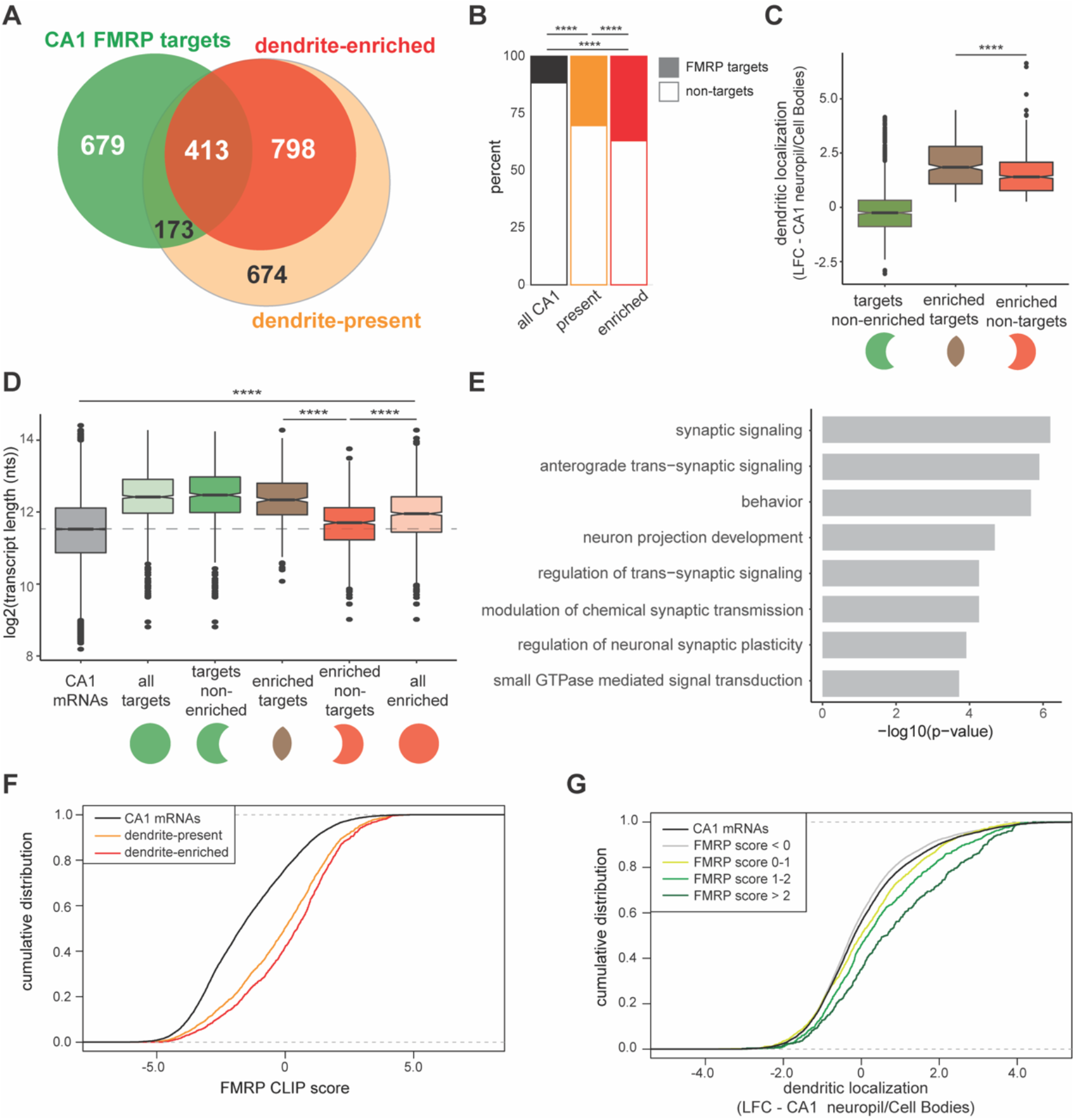
CA1 FMRP-targets are over-represented in the dendritic transcriptome. A) *Overlap of CA1 FMRP targets and the dendritic transcriptome defined CA1 neuropil TRAP.* CA1 FMRP targets are defined as those with FMRP CLIP-scores > 1 in hippocampal CamK-cTag-FMRP (Sawicka et al., 2019). B) *CA1 RP targets are more over-represented in the dendrite-enriched mRNAs than in the dendrite-present mRNAs.* Chi-squared analysis was performed to determine the enrichment of CA1 FMRP targets among dendrite-present mRNAs (p-value = 1.42e-175) and dendrite-enriched mRNAs (p-value = 1.70e-170). *CA1 FMRP targets are highly localized to the dendrites* Dendrite-enriched mRNAs were subdivided into CA1 FMRP targets and non-targets, and the dendritic localization (defined by LFC CA1 neuropil/Cell Bodies in DESeq2 analysis) was compared for each group. Dendrite-enriched mRNAs that are also CA1 FMRP gets are significantly more localized than dendritic non-FMRP targets. Wilcoxon rank sum test was used to determine significance (**** = p-value < .00001). *CA1 FMRP targets in the dendritic transcriptome are significantly longer than non-FMRP targets.* mRNA transcript lengths (in log2(nts)) for all CA1 expressed genes and the subsets defined in A were compared. For each gene expressed in the CA1 transcriptome, the length of the most highly expressed mRNA was considered. Wilcoxon rank sum test was used to determine significance. Dashed line indicates the mean transcript length for all CA1 mRNAs. E) *CA1 FMRP gets in the dendritic transcriptome encode proteins involved in synaptic signaling and synaptic plasticity*. GO analysis was performed by comparing the ndrite-enriched CA1 FMRP targets with all dendrite-enriched mRNAs. F) *CA1 FMRP targets in the dendritic transcriptome have large CA1 FMRP CLIP scores.* 1 FMRP CLIP scores for all CA1 genes were determined previously for whole-cell FMRP cTag CLIP and CA1-specific TRAP. CDF plots compare the CA1 RP CLIP scores for all CA1 genes (black) and those defined as either dendrite-present (orange) or dendrite-enriched (red). G) *Dendritic localization of CA1 RP targets found in the dendritic transcriptome correlates with FMRP binding*. Dendritic localization (LFC CA1 neuropil TRAP vs Cell Bodies TRAP) was compared by CDF plots for all CA1 genes (black) and subsets with CA1 FMRP-CLIP scores less than 0, 0-1, 1-2 or over 2.

Subdividing dendrite-enriched mRNAs into CA1 FMRP targets and non-targets revealed unique characteristics of each group. Dendrite-enriched CA1 FMRP targets were significantly more localized than dendrite-enriched non-FMRP targets in microdissected TRAP samples (Figure 4C). FMRP has been shown to preferentially bind long mRNAs (Darnell et al., 2011; Sawicka et al., 2019). While we observed that dendrite-enriched mRNAs were longer, on average, than all CA1-expressed mRNAs (Figure S2), CA1 FMRP targets in the dendrite-enriched group are significantly longer than dendrite-enriched non-targets (p-value = 9.13e-46, Figure 4D), suggesting that FMRP binds the majority of long, dendritically localized mRNAs.

Examination of the function differences between dendrite-enriched FMRP targets and non-targets revealed an enrichment in dendrite-localized FMRP targets for proteins involved in synaptic signaling, behavior, regulation of trans-synaptic signaling, and GTPase mediated signal transduction (Figure 4E), indicating that FMRP may be a key regulator of local translation of proteins involved in key synaptic functions.

FMRP CLIP scores were previously developed as a metric to define high-affinity FMRP-bound transcripts in CA1 neurons (Sawicka et al., 2019). Dendrite-enriched mRNAs had significantly higher FMRP CLIP scores than the dendrite-present group indicating greater FMRP binding (p-value = 2.646e-05, Figure 4F). Additionally, FMRP CLIP scores positively correlated with dendritic localization: when CA1 mRNAs were grouped according to the magnitude of their CA1 FMRP CLIP scores, those with increasingly higher scores were increasingly localized in the dendrites (Figure 4G). Taken together, these results demonstrate that dendrite-enriched CA1 FMRP targets constitute a highly localized subset of all dendrite-enriched mRNAs. Moreover, the magnitude of CA1 FMRP CLIP scores predict the degree of localization of its targets (Figure 4G).

### FMRP selectively binds localized mRNA isoforms

We examined whether differentially localized transcript isoforms were specifically bound by FMRP in hippocampal CA1 neurons. For example, the *Ankrd11* transcript undergoes APA to express a short and long isoform, and the long isoform is specifically localized to CA1 dendrites (Figures 2I-2K). Interestingly, CA1 FMRP-CLIP tags were detected on the long, localized isoform, but only sparsely found on the short isoform (Figure 5A-grey dashed boxes). To look at this phenomenon on a transcriptome-wide scale, we isolated exon junction reads in whole hippocampus CA1 FMRP-cTag-CLIP data. While the length of CLIP tags (20-100 nts) results in a low number of junction reads, we were able to confidently identify FMRP-CLIP tags covering 17 differentially localized alternative splice events. For example, FMRP binding was largely absent on a shorter, CB-enriched isoform of the *Cnksr2* transcript, while robust binding was evident on the longer, localized 3’UTR (Figure 5B, grey dashed boxes). Of the 12 exon-junction reads that originated from exon 20 of the *Cnksr2* transcript, 10 were derived from the long isoform, suggesting that approximately 80% of the FMRP-bound *Cnksr2* transcripts derived from the longer, dendritically localized isoform. This was especially striking since the shorter isoform was the predominant isoform in CA1 pyramidal neurons (∼80% of exon junction reads in cell body TRAP belonged to the short isoform), indicating a high degree of selectivity of FMRP binding to the dendritically localized isoform (Figure 5B). Globally, we compared the PSI values determined for the 17 detected alternative splice events detected in FMRP-CLIP with those in the CA1 cell body- and neuropil-TRAP data. This revealed that splicing events identified in FMRP bound mRNAs showed stronger correlation with PSI values determined in CA1 neuropil TRAP relative to cell body TRAP (Figure 5C). Taken together, these results indicate that FMRP preferentially binds to specific processed transcripts that are fated for dendritic localization.

**Figure 5.**
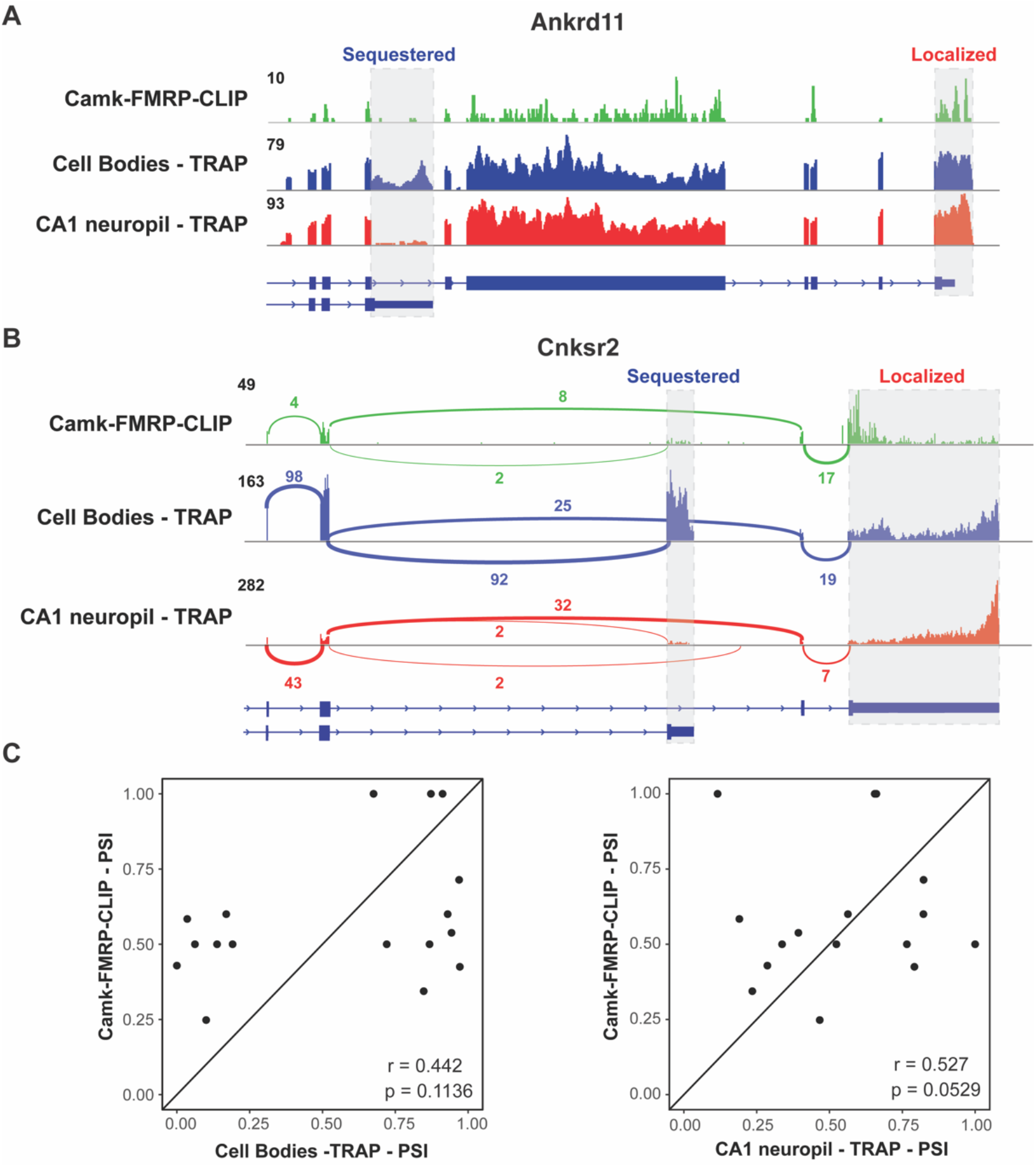
FMRP specifically binds localized mRNA isoforms. A) *FMRP preferentially binds long, localized Ankrd11 mRNAs.* The *Ankrd11* gene encodes mRNAs with two potential 3’UTRs (gray boxes). CA1 FMRP-CLIP tags (from hippocampal Camk-cTag FMRP CLIP reported previously (Sawicka et al., 2019)) are shown in green, and representative coverage from CA1 Cell Bodies- (blue) and CA1 neuropil-(red) TRAP is shown. B) *Splice junctions in FMRP-CLIP derived from Cnksr2 mRNA isoforms. Cnksr2* expresses two mRNA isoforms, indicated by gray boxes. Sashimi plots illustrate coverage and junction-spanning reads from CA1 FMRP-CLIP in green (tags are aggregated from three replicates). Sashimi plots are also shown for Cell Bodies TRAP (blue) and CA1 neuropil-TRAP (red). C) *Splicing isoforms discovered in FMRP-CLIP tags resemble those found in the localized transcriptome.* PSI values derived from splice junction reads in CA1 FMRP-cTag-CLIP tags were compared to PSI values from the same events in Cell Bodies-TRAP (left) and CA1 neuropil TRAP (right). Results of Pearson correlation test is shown.

### Identification of dendritic FMRP targets

In order to identify direct FMRP-bound mRNA targets in CA1 dendrites, we crossed FMRP cTag mice with CamkIIα-Cre mice, tagging FMRP with GFP specifically in the CA1 pyramidal neurons (Figure 1A). Hippocampal slices from cTag mice were crosslinked, microdissected into cell body and neuropil regions, and subjected to FMRP-CLIP using antibodies against GFP. This allowed purification of FMRP-bound RNA specifically in the CA1 cell bodies or dendrites. Across five biological replicates, we obtained 746,827 FMRP CA1-specific CLIP tags from the cell bodies and 80,749 tags from CA1 dendrites. Overall, we observed a similar distribution of FMRP CLIP tags across the CDS in these mRNAs and in the two compartments, consistent with prior CLIP analysis and the general observation that FMRP binds CDS to arrest ribosomal elongation ((Darnell et al., 2011); Figure S5).

Combining compartment-specific TRAP and FMRP-CLIP experiments allowed us to determine compartment-specific FMRP CLIP scores for the CA1 cell bodies and dendrites (Figure 6A, Figure S6, Supplemental Table 7). From this, we identified 383 mRNAs which are reproducibly bound by FMRP in CA1 dendrites (Supplemental Table 8). Of these dendritic FMRP targets, 60.8% (233) were dendrite-enriched (Figure 6B), and 76.5% (293) were dendrite-present (Figure S5). As anticipated, dendritic FMRP targets show greater dendritic localization in TRAP when compared to all CA1 FMRP targets (Figure 6C). Additionally, when considering the FMRP-CLIP scores identified previously by whole hippocampus CA1 FMRP-CLIP, the scores for the dendritic FMRP targets were significantly larger than the scores for all dendrite-enriched mRNAs (Figure 6D), suggesting that the dendritic FMRP targets identified here represent a subset of previously-identified FMRP targets for which high-affinity FMRP binding is consistent with significant localization into the CA1 dendrites.

**Figure 6.**
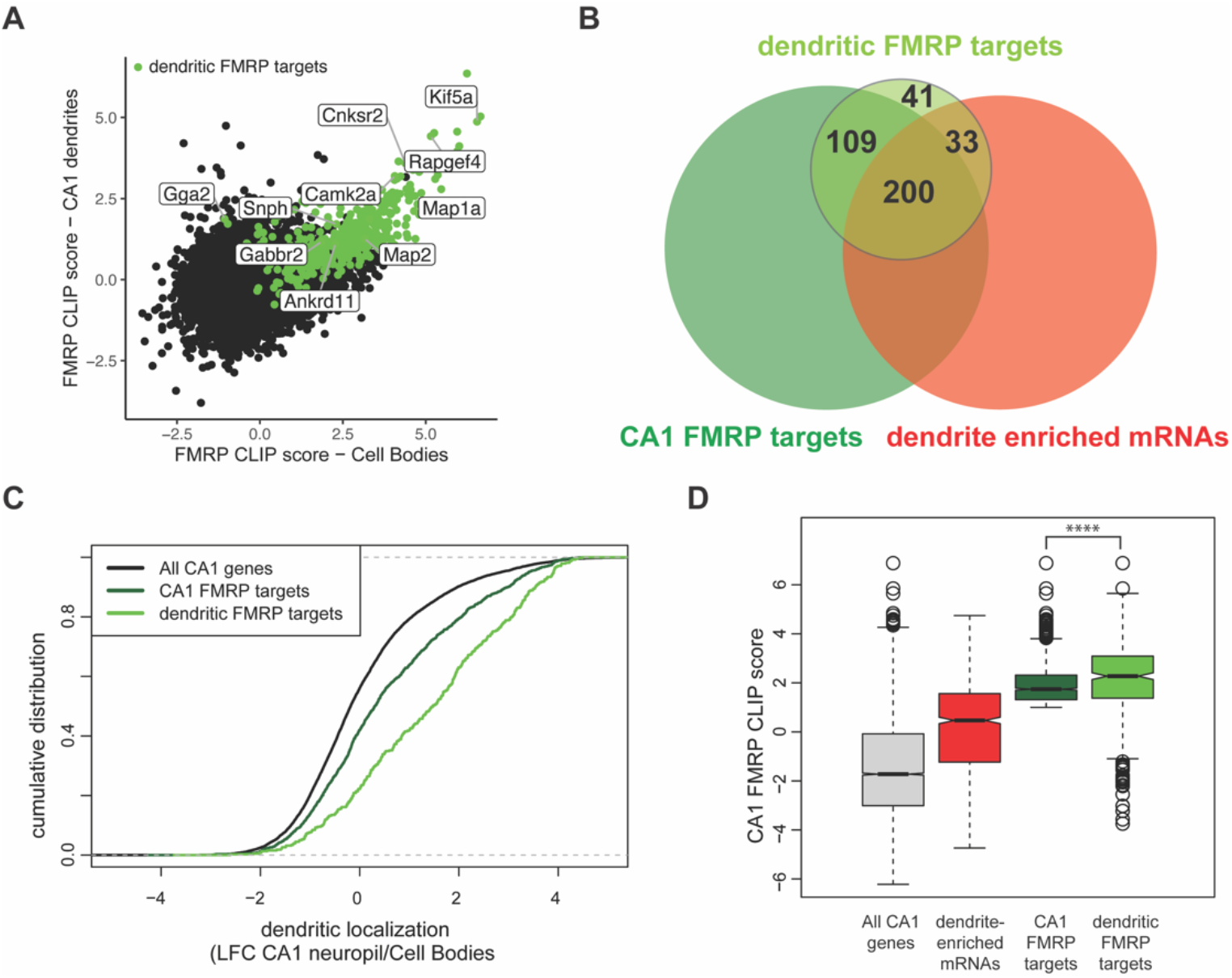
Compartment-specific cTag FMRP-CLIP reveals dendritic FMRP-targets. A) *Compartment-specific Camk-cTag FMRP-CLIP and TRAP-seq were integrated to determine compartment-specific FMRP CLIP scores*. CLIP scores were determined for all replicates. Plotted is the mean CLIP scores for the CA1 Cell Bodies and dendrites. Dendritic FMRP targets are colored in green. Genes of interest are labeled. B) *A subset of CA1 FMRP-CLIP targets are also dendritic FMRP-CLIP targets.* These are overlapped with dendrite-enriched mRNAs and whole-cell CA1 FMRP targets. C) *Dendritic FMRP targets are highly localized.* Dendrite-enrichment (LFC CA1 neuropil-TRAP/Cell Bodies-TRAP) is plotted for all CA1 genes, all CA1 FMRP targets, and dendritic FMRP-CLIP targets. D) *Dendritic FMRP targets have high FMRP binding scores.* Whole-cell CA1 FMRP CLIP scores are plotted for all CA1 mRNAs, dendrite-enriched mRNAs, all CA1 FMRP targets and dendritic FMRP targets. Stars indicate significance in Wilcoxon rank tests (**** = p-value < .00001).

### Subcellular compartment-specific FMRP-CLIP scores reveal functionally distinct groups of FMRP targets

Many directly-bound FMRP target transcripts encode proteins that are implicated in Autism Spectrum Disorders (ASD) (Darnell et al., 2011; Iossifov et al., 2012; Zhou et al., 2019). We hypothesized that FMRP may regulate functional subsets of its targets in a subcellular-compartment specific manner, a phenomenon that would be reflected by differences in compartment-specific FMRP binding. To test this, we segregated all whole-cell CA1 FMRP CLIP targets according to their function by module detection using the HumanBase software (Krishnan et al., 2016). Eight functional modules were detected, three of which contain more than 100 genes (Figure 7A, Supplemental Table 9). The FM1 cluster, which contains 393 genes, is highly enriched for genes involved in nuclear regulation of gene expression, with the top GO terms being chromatin organization and modification and histone modification. FM2 (292 genes) is enriched for genes involved in ion transport and receptor signaling. The FM3 cluster (203 genes) contains genes involved in maintenance of cell polarity and autophagy (Figure 7B).

**Figure 7.**
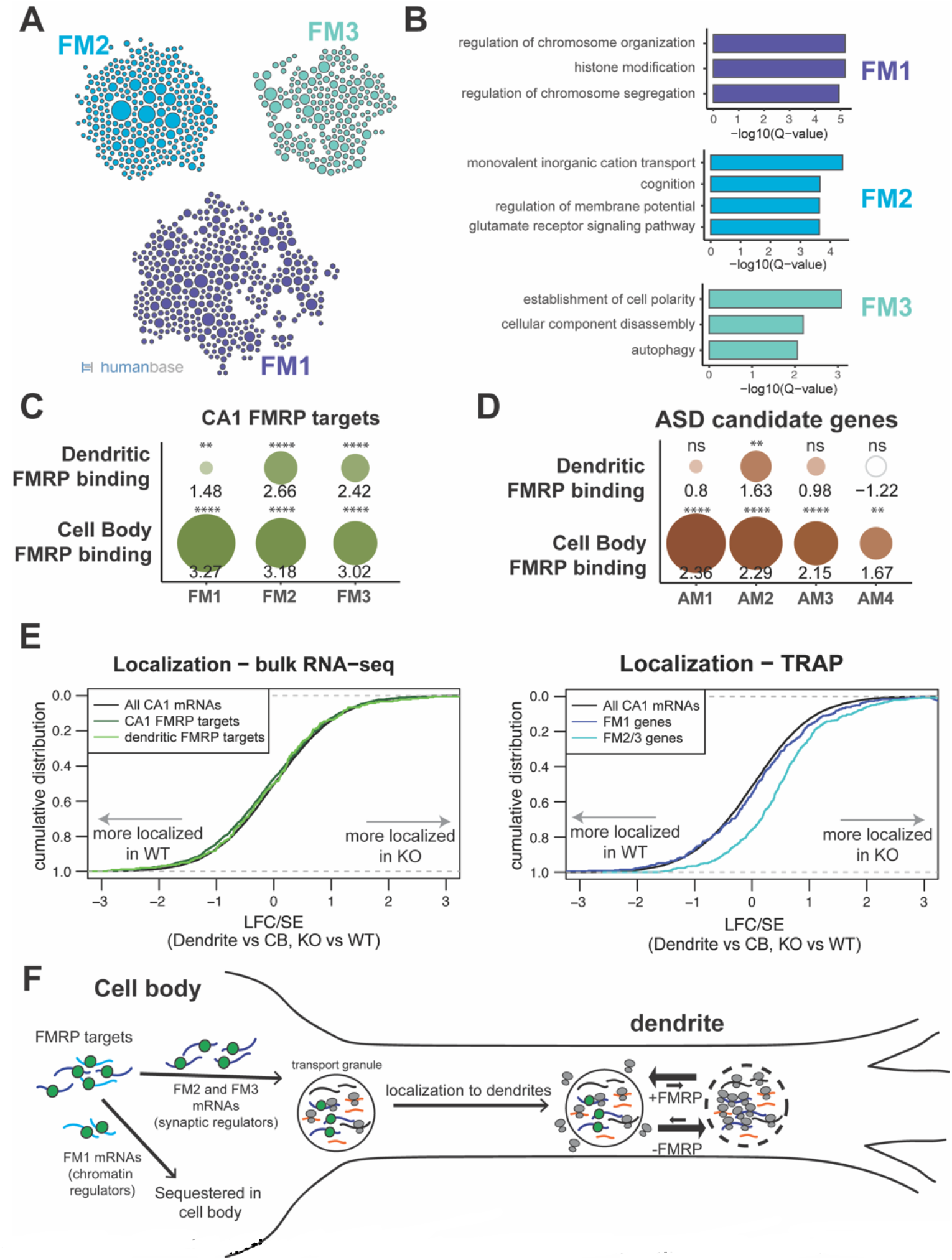
FMRP regulates functionally distinct mRNAs in the cell bodies and dendrites of CA1 neurons. A) *CA1 FMRP targets fall into three functional clusters.* Functional module detection was performed for CA1 FMRP targets by the HumanBase software. B) *Top GO terms for the three largest functional modules of CA1 FMRP targets.* Q-values for enrichment of terms were determined by the HumanBase software. C) *Dendritic FMRP targets are enriched in functionally distinct modules of CA1 FMRP targets.* CA1 genes were ranked according to FMRP dendritic and cell bodies FMRP-CLIP scores and GSEA analysis was performed using the FMRP functional clusters (from *A*) as gene sets. Circles are colored according to normalized enrichment scores (NES) and sized according to FDR from the GSEA analysis. NES values are shown, and stars indicate significance (** is FDR < .001, **** is FDR < .00001). D) *Dendritic FMRP targets are enriched in a functional module of autism candidate genes.* GSEA analysis was performed as shown in *C*, with functional modules of autism candidate genes (SFARI) clustered according to the HumanBase software. E) *Localization of FM2/3 FMRP targets are largely unchanged in compartment-specific bulk RNA-seq of FMRP* KO *animals, but increased in TRAP.* (left) Neuropil localization (LFC/SE of CA1 neuropil bulk RNA-seq vs Cell Bodies bulk RNA-seq) was assessed in FMRP KO vs WT animals. Cumulative distribution plots are shown. Shifts to the right indicate more localization in the FMRP KO animals, and shifts to the left indicate more localization in WT animals. All CA1-expressed genes, all CA1 FMRP targets and dendritic FMRP targets are shown. (right) Neuropil localization in TRAP-seq on FMRP KO vs WT animals, with subsets including the FM1/2/3 groups of CA1 FMRP targets as described in A. F) *FMRP regulates functionally distinct subsets of mRNAs in CA1 cell bodies and dendrites.* FMRP binds its target mRNAs in the cell bodies, including both FM1 (chromatin regulators) and FM2/3 (synaptic regulators). FM1 mRNAs are sequestered in the cell bodies and FM2/3 mRNAs are transported to CA1 dendrites, where they are subject to FMRP regulation, which alters the degree of ribosome binding. In FMRP containing neurons, target mRNAs are less ribosome bound than in FMRP-deficient neurons.

In order to determine if any of these functional modules might be differentially regulated by FMRP in the dendrites and cell bodies of CA1 neurons, we performed GSEA analysis to estimate enrichment of the FM1-3 transcripts among all FMRP-bound, CA1-expressed transcripts ranked by their dendritic or cell bodies-specific FMRP CLIP score. We found that while the FM2 and FM3 clusters were highly enriched in FMRP-bound mRNAs in both the dendrites and the cell bodies, the FM1 cluster was strongly enriched among cell body-bound FMRP targets, but only weakly enriched among the dendritic FMRP bound transcripts (Figure 7C). This suggests that FM2 and FM3 modules contain mRNAs that are directly bound and regulated by FMRP in dendrites, while the FM1 cluster contains highly bound FMRP targets in the cell bodies, indicating distinct, biologically coherent regulation.

We further utilized compartment-specific FMRP-CLIP scores to identify functional modules of ASD candidate mRNAs subject to compartment-specific FMRP regulation (Figure 7D and S7A-S7B). One module, AM2, contains transcripts enriched for glutamate signaling, learning and memory, and is bound by FMRP in both the dendrites and cell bodies. The AM1 module consists of genes involved in chromatin modification and is highly enriched among mRNAs bound by FMRP in the cell bodies, but is not significantly enriched among dendritic FMRP-bound mRNAs. This suggests the possibility of compartmentalized roles for FMRP in which mRNAs important for synaptic signaling are bound and regulated by FMRP near the synapses, while mRNAs bound by FMRP in the cell bodies are involved in regulation of neuronal gene expression through chromatin regulation.

### Dysregulation of mRNA localization in FMRP KO animals

In order to look for FMRP-dependent regulation of dendritic mRNAs, we performed bulk RNA-seq and cell-type specific TRAP on microdissected hippocampi from WT and FMRP KO mice. Bulk RNA-seq of microdissected material in FMRP WT and KO mice showed no overall change in the localization of FMRP targets (Figure 7E, left panel). In contrast, TRAP revealed that mRNA levels of ribosome-associated FMRP targets were increased in CA1 dendrites of KO mice. Interestingly, this was only evident for FM2/3 genes, not FM1 genes (Figure 7E, right panel). While FMRP targets are generally downregulated in TRAP from hippocampal neurons (Sawicka et al., 2019), a finding that we replicate in cell bodies, transcripts that encode synaptic regulatory proteins (FM2/3), which are bound by FMRP in the dendrites, show increased ribosome association in CA1 dendrites of KO animals (Figure S7C). These surprising results suggest a model in which FMRP may be differentially involved in translational regulation of functionally distinct mRNAs in specific neuronal compartments. (Figure 7F).

## Discussion

Recent advances in cell-type specific transcriptomic approaches have greatly increased the resolution at which we understand gene expression in the nervous system. Here we build on these advances by incorporating compartment-specific CLIP and TRAP in order to define a high-quality, cell-type specific transcriptome of CA1 neuronal cell bodies and dendrites *in vivo*, and to define compartment-specific FMRP regulation of its targets. We found subcellular differences in the sets of alternatively spliced or polyadenylated transcripts in each compartment, connecting pre-mRNA nuclear regulation to subcellular localization in neurons (Figures 2 and 3). Moreover, previously-defined FMRP targets are overrepresented in the dendritic transcriptome, and FMRP often preferentially binds localized mRNA isoforms. We define dendritic FMRP targets and find that their dendritic ribosome association is increased in FMRP-null mice, consistent with differential translational regulation between subcellular compartments. Distinct sets of FMRP-bound autism-related transcripts have been described--particularly those related to chromatin regulation and those related to synaptic plasticity (Darnell et al., 2011; Iossifov et al., 2012). Remarkably, we find here that these transcripts show different subcellular localization--transcripts encoding chromatin regulators are enriched in CA1 cell bodies, while those encoding synaptic regulators are enriched in their dendrites. Together, these observations indicate that RNA regulatory factors link post-transcriptional controls with subcellular localization of RNA isoforms in neurons. The data support and extend a model (Darnell, 2020) in which FMRP integrates cellular activity and signaling to maintain neuronal homeostatic plasticity (Turrigiano, 2012) by mediating differential translation of transcripts encoding nuclear and synaptic functions in the cell body and and dendrite, respectively (Figure 7F).

Although RNA-sequencing (Cajigas et al., 2012), 3-Seq (Tushev et al., 2018) and TRAP-seq (Ainsley et al., 2014) have been performed previously for microdissected CA1 neuropil, these studies were either not performed using cell-type specific approaches or unable to capture full-length mRNAs. As the mRNAs presented here are intact and relatively free of contaminating cell-types (Figure S1), this dataset can be used for definition of dendritic mRNAs and for identification of differential localization of 3’UTR (Figure 2) and alternative splice isoforms (Figure 3), making it a valuable dataset for the community as a whole.

Consistent with prior reports (Tushev et al., 2018), we find that the majority of differentially localized 3’UTRs are longer than their sequestered counterparts, suggesting that alternative polyadenylation events that lead to longer 3’UTR isoforms could lead to the inclusion of regulatory elements that allow for localization, such as binding sites for RNA-binding proteins or Ago-miRNA complexes. Long 3’UTRs may also act to recruit binding partners for nascent proteins, which can affect the function and/or localization of the protein, as previously reported (Berkovits and Mayr, 2015). Future experiments analyzing compartment-specific cTag-CLIP of RNA-binding proteins that bind to 3’UTRs such as Ago, Staufen, Nova1/2 or Elavl3/4 will provide further insight into the role of these 3’UTRs in mRNA localization and local translation. In addition to 3’UTR-APA, 20% of differentially localized 3’UTRs result from APA events which impact the coding sequence, indicating that differential mRNA localization may lead to expression of functionally distinct protein isoforms in the two compartments.

The quality and purity of our data also allowed for detection of alternative splicing events that result in differentially localized mRNA alternatively spliced isoforms, which has not been reported previously. Interestingly, we found that NOVA2, a neuron-specific alternative splicing factor, is responsible for the generation of splicing isoforms that are sequestered to the neuronal cell bodies of CA1 neurons (Figure 3E). NOVA1 is one of a relatively small number of mammalian splicing factors demonstrated to directly bind to pre-mRNA and thereby regulate alternative splicing (Licatalosi et al., 2008; Zhang and Darnell, 2011) and to bind 3’ UTRs of those same transcripts (Eom et al., 2013), and in the case of GlyRa2, to co-localize with the same transcript in dendrites (Racca et al., 2010). These findings further underscore the many ways in which RNA-binding proteins contribute to neuronal complexity in specific subcellular compartments.

The significant overlap between CA1 FMRP targets and dendrite-enriched mRNAs supports literature indicating that FMRP is an important regulator of the dendritic transcriptome (Bagni and Zukin, 2019; Banerjee et al., 2018; Liu-Yesucevitz et al., 2011). Further, overall FMRP binding affinity (defined by FMRP-CLIP scores in hippocampal neurons) correlates with the degree of dendritic localization of a given mRNA (i.e. the enrichment of mRNAs in the neuropil over the cell body, Figure 4G), indicating a strong relationship between FMRP binding affinity and mRNA localization.

Interestingly, through analysis of whole-cell cTag FMRP-CLIP data, we find multiple instances of FMRP selectively binding to localized isoforms (Figure 5). A striking example is the case of the *Cnksr2* gene, which generates a short, sequestered mRNA and a longer, highly localized isoform. The protein encoded from the localized mRNAs contains an additional PDZ-binding domain that is not present in the shorter isoform. *Cnksr2* has been identified in GWAS studies as an ASD candidate, and mutations in this gene have been shown to cause epilepsy and intellectual disability (Aypar et al., 2015). In the cell body compartment, the shorter isoform is predominant, which can be seen by both PAPERCLIP and TRAP. However, FMRP-CLIP, which generally binds the CDS and at least the proximal 3’UTR regions of its targets, shows predominant binding on the 3’UTR of a minor isoform in the presence of a more highly expressed, shorter, dendrite-localized mRNA isoform (Figure 5B). Similar trends can be seen with a number of other mRNAs such as *Ankrd11* (Figure 5) and *Anks1b* (data not shown). Taken together, this data indicates that FMRP can display binding preferences both on different transcripts and different isoforms of the same transcript. This adds an additional layer to the already-complicated process of how FMRP recognizes and binds its targets, and may suggest that FMRP binding specificity may rely on localization-determining events in the nucleus, such as deposition of RNA-binding proteins on the 3’UTRs of alternatively spliced transcripts.

FMRP binding to localized mRNA isoforms may also be a result of events that occur in the cytoplasm. For example, some mRNAs with longer 3’UTRs themselves may possess great propensity for entrance into FMRP-containing transport granules due simply to length, similar to the findings that long mRNAs are preferentially found in stress granules due to lower translation efficiency and increased ability for RNA-RNA interactions to form, which are thought to stabilize RNA granules (Khong et al., 2017). This is supported by findings that FMRP is found in neuronal mRNA transport granules (Dictenberg et al., 2008) and is also known to bind to RNA structural elements such as kissing complexes and G-quadruplexes (Darnell et al., 2005), and suggests a role for FMRP in maintaining the translationally repressed status of long mRNAs in transport granules.

We present here the first reported list of mRNAs that are highly bound to FMRP in the dendrites of CA1 neurons. These 383 targets are highly enriched in the dendrites and show high affinity FMRP binding in whole CA1 neurons (Figure 6). These targets were determined by combining compartment-specific CLIP and TRAP experiments to determine compartment-specific FMRP CLIP scores. Importantly, these targets are the result of stringent filtering to include only high-confidence dendritic FMRP targets. However, we bioinformatically extended our findings using compartment-specific FMRP CLIP scores to identify functional clusters of FMRP targets that are differentially localized. We find a remarkable link between the function of a given FMRP target and its subcellular localization. FM1, which contains transcripts encoding proteins with nuclear functions such as histone modification and chromosome organization are enriched in CA1 cell bodies, while the FM2 and FM3 mRNAs, which encode for proteins with synaptic functions such as ion transport and cell polarity are found in both cell bodies and dendrites.

Interestingly, mRNAs from genes in the FM2 and FM3 clusters show increased ribosome association in the FMRP KO mouse in a pattern distinct from the FM1 genes (Figure 7E). However, bulk RNA-seq on the same compartments showed that overall FMRP targets levels were largely unchanged in the neuropil. This suggests that while FMRP targets can be localized to dendrites in the absence of FMRP, they may be increasingly ribosome associated there. This supports the proposal (Wang et al., 2008) that FMRP in the neuronal processes may exist in a polyribosome-depleted granule which is altered to become translationally competent upon neuronal activity. It is also consistent with the detection of increased basal translation rates in mouse models of Fragile X Syndrome (Gross et al., 2010; Liu et al., 2012). Taken together, we suggest a model in which FMRP specifically binds mRNAs encoding synaptic proteins and fated for dendritic localization, and maintains them in a translationally repressed, and potentially polyribosome-depleted, state for transport into the processes. Further, within the dendrite, our findings in FMRP-null mice are consistent with a role in the ability of neuronal activity to induce polyribosome formation and local translation (and concomitant increased polyribosome density) of its specific targets (Figure 7F). Future experiments investigating how dendritic FMRP binding changes upon neuronal activity will help to elucidate the precise role of FMRP in regulation of activity-dependent local translation.

In summary, we present data demonstrating the ability to utilize compartment- and cell-type specific profiling technologies to precisely define the dendritic transcriptome. Our results underscore the role of FMRP as an important regulator of the dendritic transcriptome, playing an important role in the ribosome-association of isoform-specific dendritic targets. This finding, coupled with the identification of the FM1 genes that are regulated by FMRP exclusively in the cell bodies supports the hypothesis (Darnell, 2020) that FMRP acts as a sensor for neuronal activity through actions in both the cell bodies and dendrites of neurons. Further studies into how these groups of genes are differentially FMRP-regulated in subcellular-compartment specific manner will have important implications in the understanding of how dysregulation of FMRP and its targets lead to intellectual disability and ASD.

## Methods

### Mice

All mouse procedures were conducted according to the Institutional Animal Care and Use Committee (IACUC) guidelines at the Rockefeller University. RiboTag (B6N.129-Rpl22^tm1.1Psam^/J, stock no. 011029) and Camk2a-Cre (B6.Cg-Tg(Camk2a-cre)T29-1Stl/J, stock no. 005359) were obtained from Jackson Laboratories. FMRP cTag (Van Driesche et al, 2019) and Pabpc1 cTag (Hwang et al., 2017) mice were previously described. B6.129P2-Fmr1tm1Cgr/J (Fmr1 KO) mice were a generous gift from W.T. Greenough maintained for multiple generations in our own facilities. Mice were housed up to 5 mice per cage in a 12 hr light/dark cycle. Breeding schemes for TRAP-seq (producing RiboTag^+/-^, FMRP^+/+^ and RiboTag^+/-^,FMRP^Y/-^ male littermates) and FMRP cTag-CLIP (producing Cre^+/-^; Fmr1-cTag^+/Y^ male offspring) were described previously (Sawicka et al., 2019).

### Immunofluorescence

Immunoflourescence was performed as described previously (Sawicka et al., 2019). Primary antibodies used were NeuN (Millipore ABN90P, RRID:AB_2341095, 1:2000 dilution), and HA (Cell Signaling, C29F4, RRID:AB_1549585, 1:4000 dilution).

### TRAP- and RNA-seq of microdissected hippocampal slices

For each TRAP-seq replicate (four replicates were performed), hippocampi from three adult mice (6-10 weeks) were sectioned into 300 μm slices using a tissue chopper and microdissected in HBSS containing 0.1 mg/mL cycloheximide. For microdissection, the CA1 was excised from the hippocampal slices and separated into a cell body (CB) and neuropil layer. Microdissected tissue from each mouse was collected and resuspended in 0.5 mL ice-cold polysome buffer (20 mM Hepes, pH 7.4, 150 mM NaCl, 5 mM MgCl_2_, 0.5 mM DTT, 0.1 mg/mL cycloheximide) supplemented with 40 U/ml RNasin Plus (Promega) and cOmplete Mini EDTA-free Protease Inhibitor (Roche) and homogenized by mechanical homogenization with 10 strokes at 900 rpm. NP-40 was added to 1% final concentration and incubated on ice for 10 minutes. Samples were pooled and centrifuged at 2000 x g for 10 minutes. Supernatant was subsequently centrifuged at 20,000g for 10 minutes. 10% of the resulting lysate was used for RNA-seq, and the remaining lysate was subject to pre-clearing with 1.5 mg (50 ul) Protein G Dynabeads for 45 minutes. HA-tagged ribosomes were collected by indirect IP by adding 40 ug of anti-HA antibody (Abcam ab9110, RRID:AB_307019) to CB lysate pools and 5 ug to NP lysate pools. Immunoprecipitation was incubated overnight with rotation at 4°C. Antibody-ribosome complexes were collected by addition of 7.2 mg (CB pools) or 4.44 mg (NP pools) Protein G Dynabeads and further incubated with rotation at 4°C for 1 hour. Beads were washed with 1 mL polysome buffer containing 1% NP-40 once for 5 minutes and twice for 20 minutes, followed by 4 x 10 minute washes in 50 mM Tris pH 7.5, 500 mM KCl, 12 mM MgCl2, 1% NP-40, 1 mM DTT, 0.1 mg/mL cycloheximide. RNA was extracted from beads by incubating in 500 uL Trizol at room temperature for 5 minutes. RNA was collected by standard Trizol (Invitrogen) extraction via manufacturer’s protocol, and quantified with RiboGreen Quant-IT assays (Invitrogen). Bulk RNA-seq samples were treated with RQ1 RNase-free DNase (Promega) prior to library preparation. RNA was further purified for polyadenylated RNA by using Dynabeads mRNA Purification Kit (Ambion). The libraries were prepared by TruSeq RNA Sample Preparation Kit v2 (Illumina) following manufacturer’s instructions. High-throughput sequencing was performed on HiSeq (Illumina) to obtain 100 nucleotide paired-end reads.

### Fluorescence in situ hybridization (FISH) with RNAscope

Mice were anesthetized with isoflurane and transcardially perfused with PBS containing 10 U/ml heparin followed by perfusion with ice-cold PBS containing 4% paraformaldehyde. After perfusion, animals were decapitated, and intact brains removed and postfixed overnight in 4% paraformaldehyde in PBS at 4°C. Brains were then transferred to PBS with 15% sucrose for 24 hr followed by PBS with 30% sucrose for a further 24 hr and then embedded and frozen in OCT medium. 12 μm coronal slices were prepared using a Leica CM3050 S cryostat and directly adhered to Fisherbrand 1.0mm superfrost slides (Cat. No. 12-550-15) and stored at -80C until use. FISH was performed using the RNAscope Multiplex Fluorescent Kit v2 as recommended for fixed frozen tissue, with some exceptions. For pretreatment of samples prior to hybridization, slides were baked at 60°C for 45 minutes, followed by fixation in 4% paraformaldehyde in PBS at 4°C for 90 minutes. Samples were dehydrated in ethanol [50%, 70%, 100% twice each] and incubated at room temperature before hydrogen peroxide treatment for 10-20 minutes, followed by target retrieval as recommended. After probe hybridization, samples were washed three times for 15 minutes in wash buffer heated to 37°C. Probes used were conjugated with Alexa fluorescein (488nm), Alexa Cyanine 3 (555nm), Alexa Cyanine5 (647nm). RNAscope probes were designed to recognize unique 3’UTR sequences (for UR-APA events) or for common and distal 3’UTRs (for UTR-APA events) with at least 500-1000 nts between regions. See Supplemental Table 2. Each FISH experiment was performed on at least three slices from at least two different mice.

### Image processing and quantitation

Airyscan-Fast (AS-F) image capturing was performed using the Zen Black 2.3 SP1 FP3 acquisition software on an Inverted LSM 880 Airyscan NLO laser scanning confocal Microscope (Zeiss) outfitted with AS-F module (16 detectors) and argon laser for 488 line. Objective: Zeiss Plan 63x 1.4 NA Apochromat oil immersion; imaging at this objective was performed using Immersol 518 F immersion media (ne = 1.518 (23 °C); Carl Zeiss). Acquisition parameters include laser lines: 405nm, 488nm, 561nm, 633nm [laser power adjusted until relative power for each line eliminates as much background as possible without diminishing signal]. Emission filter for Airyscan detection: 405ch, BP 420 - 480 + BP 495-620; 488ch, BP 420-480 + 495-550; 561ch, BP 420-480 + 495-620; 633ch, BP 570-620 + LP645. Settings: 8 bit-depth and acquired with image size: 135.0 x 135.0 um; Pixel size: 0.14um (step size is 0.159 using a piezo stage). All raw image data was sent directly to ZEN 2.3 software for reconstruction. Files underwent Airyscan processing (Parameters: auto strength at 6 for 3D images) before being stitched at a normalized cross-correlation threshold set at 7. Processed and stitched .czi files were converted to .ims files using Imaris File Converter x64 9.6.0 before being uploaded into Imaris x64 9.6. Spots were quantified using the spot counting operation (Imaris software) with the default values and modifying the spot detection parameters (“Model PSF-elongation along Z-axis”: Estimated XY Diameter: 0.8μm; Estimated Z Diameter: 1.4μm). Detection threshold was adjusted manually until all false/weak signals were eliminated. The mRNA coordinates (X, Y, Z) were downloaded for bioinformatic analysis. Max projections exported from Imaris were uploaded in Fiji. Images were adjusted to 8-bit, orientation is adjusted and channels are separated. For detection of nuclei for bioinformatic analysis, threshold was adjusted until the majority of the DAPI stain was detected and applied. ‘Analyze particles’ operation was applied with the settings: size 50-infinity (pixel units); circularity 0.0-1.0; show ‘masks’. Resulting text image files were used for downstream analysis.

### Compartment-specific cTag FMRP-CLIP

Microdissection of hippocampal slices from 5 - 8 adult Camk-FMRP-cTag mice was performed as described above, except that the slices were UV crosslinked in HBSS with 0.1 mg/mL cycloheximide three times using 400 mJ/cm^2^ after sectioning and before microdissection. After dissection, samples were collected and homogenized in lysis buffer (1x PBS, 0.1% SDS, 0.5% NP-40, 0.5% Sodium deoxycholate supplemented, 1X cOmplete Mini EDTA-free Protease Inhibitor (Roche) and 0.1 mg/mL cycloheximide) by passing through syringes with a 28 gauge needle. cTag FMRP-CLIP was performed as described previously (Sawicka et al., 2019), with minor modifications. Cell body pools were lysed in 1 mL of lysis buffer and neuropil pools in 0.5 mL. Pre-clearing was performed with 6 and 1.5 mg of protein G Dynabeads for CB and NP pools, respectively. Immunoprecipitation was performed using mouse monoclonal anti-GFP antibodies conjugated to protein G Dynabeads, using 25 ug of each antibody for CB pools and 6.25 ug of each antibody for NP pools and rotated at 4°C for 1-2 hours. IPs washes were rotated 2-3 minutes at room temperature. RNA tags were cloned as described previously (Sawicka et al., 2019), with cell bodies and neuropil samples being pooled after barcoding in order to increase yield for low-input samples.

### Compartment-specific cTag PAPERCLIP

Collection and UV crosslinking of microdissected material was performed as described for compartment-specific cTag FMRP-CLIP. cTag-PAPERCLIP was performed as described previously (Hwang et al., 2017) with the following exceptions. Four replicates were performed, using 3-14 mice per replicate. CB pools were lysed in 1 mL of lysis buffer, NP pools in 0.5 mL. Additional IP washes were performed using stringent washes conditions (described in (Sawicka et al., 2019)), and low-input samples were pooled after barcoding. Cell body pools were lysed in 1 mL of lysis buffer and neuropil pools in 0.5 mL. Immunoprecipitation was performed using mouse monoclonal anti-GFP antibodies conjugated to protein G Dynabeads, using 25 ug of each antibody for CB pools and 6.25 ug of each antibody for NP pools and rotated at 4°C for 3-4 hours. RNA tags were cloned as described previously (Hwang et al., 2017) with cell bodies and neuropil samples being pooled after barcoding in order to increase yield for low-input samples

### Bioinformatics

#### Calling localized mRNAs

Transcript expression was quantified from RNA-seq and TRAP-seq using salmon and mm10 gene models. Pairwise comparisons with batch correction were performed using DESeq2 for CA1 neuropil vs cell bodies, with and without Cre expression, and TRAP vs bulk RNA-seq. Dendrite-localized genes were defined as those with a Benjamini–Hochberg FDR less than 0.05 for FDR for TRAP vs RNA-seq, log2 fold change (LFC) TRAP vs RNA-seq greater than 0, and log2 fold change Cre-positive vs Cre-negative greater than 0 (all in CA1 neuropil samples only). Dendrite-enriched mRNAs used the same filters, but also required an FDR of CA1 neuropil vs cell bodies of less than 0.05. Dendritic localization is defined as the log2 fold change resulting from DESeq2 analysis of CA1 neuropil vs Cell Bodies TRAP samples. For length analysis, the transcript that showed the highest expression in whole-cell hippocampal Camk2a-TRAP (Sawicka et al., 2019) was used.

#### FISH quantification

Nuclei (from DAPI stains) and spots (from FISH) were identified and their locations in the image determined with Fiji and Imaris software. For prediction of the location of the cell body layer in each image, nuclei and spot-containing pixels were identified and converted into scatterplots in R. Scatterplots were sliced into 25 vertical slices, and the density of each slice was plotted in order to identify the location of the bottom of the cell body layer in each slice. These points were subject to two rounds of polynomial curve fitting, with outliers removed manually between the two rounds. The predicted distance between each FISH spot and the cell body was determined using the distance between the spot and the fitted curve. For t-tests, spots were considered to be in the neuropil if they were more than 10 microns from the predicted line. Changes in distribution were also assessed using Kolmogorov–Smirnov tests.

#### Identification of differentially localized 3’UTR isoforms

polyA sites were identified from PAPERCLIP data using the CTK package (Shah et al., 2017) as described previously. From whole-cell PAPERCLIP datasets, peaks were considered that had 10 or more tags and represented 5% or more of the tags on that gene. For microdissected PAPERCLIP datasets, any peaks that had tags in more than one neuropil PAPERCLIP experiment were considered. Splice junctions were identified in both whole-cell and micro-dissected TRAP samples. Splice junctions were considered if they were found in 10 reads or if they represented 10% of total junction reads for that gene. Using the GenomicRanges package (Lawrence et al., 2013), the upstream splice junction was identified for each PAPERCLIP site, and the downstream PAPERCLIP site was identified for each splice junction. Percentage of covered bases for these potential 3’UTRs was determined using bedtools (Quinlan and Hall, 2010) and only those with 80% coverage in any single experiment were considered in downstream analyses. Next, ambiguous genes and 3’UTRs that overlapped other genes/UTR were eliminated s. This yields all expressed final exons. Genes with multiple 3’UTRs were selected and used for counting of reads from microdissected TRAP-seq samples using featureCounts, followed by DEXSeq analysis (Anders et al., 2012) to identify differentially localized 3’UTRs.

#### Splicing

Splicing analysis was performed using rMATs (Shen et al., 2014), considering both junction counts and exon coverage and the maser R package was used for visualization. For splicing analysis of RNABP KO mice, rMATs analysis was performed on datasets shown in Supplemental Table 4. Sashimi plots were generated in IGV.

#### Compartment-specific CLIP

CLIP tags were processed as described previously ((Sawicka et al., 2019) for FMRP-CLIP and (Hwang et al., 2017) for cTag-PAPERCLIP). Briefly, for FMRP-CLIP, tags were mapped to the transcriptome, using the transcript with the highest-expression for each gene as determined by whole-cell Camk2a-TRAP (Sawicka et al., 2019) . For cTag-PAPERCLIP, tags were mapped to the genome and polyadenylation sites were determined by clusters called using the CTK software (Shah et al., 2017).

#### Calling dendritic FMRP targets

Counts of FMRP-CLIP tags mapped to transcripts were normalized first for transcript length and then by sequencing depth (scaled to 10,000 tags) in order to generate length and library size normalized CLIP expression values for each transcript. mRNAs were determined to be dendritic FMRP targets if they fit one of two criteria: 1) if they were reproducibly detected in cTag-FMRP-CLIP on the neuropil (greater than 5 normalized tags per 10,000 in at least 3 of 5 replicates, 287 genes) or 2) if they had a mean compartment-specific CLIP score > 1 (241 genes). See Supplemental Table 6 for CLIP scores and CLIP expression information. CLIP scores were determined as described previously (Sawicka et al., 2019), with a few exceptions to account for low numbers of dendritic CLIP tags. All CLIP tags that map along the length of CA1 mRNAs were used for analysis. CLIP expression scores were calculated by dividing CLIP tags by transcript length, followed by normalization for library depth. TPMs for TRAP-seq were determined by the tximport package from pseudocounts obtained from salmon (Patro et al., 2017; Soneson et al., 2015). For each CLIP replicate and compartment, TRAP TPMs were plotted against CLIP expression scores with a TRAP TPM > 1 and FMRP-CLIP tags in 3 or more replicates. Linear models were determined and mean CLIP scores were calculated as described previously (Sawicka et al., 2019).

#### Functional clustering of FMRP targets

Functional Module Detection implemented within the HumanBase software was used to determine functional clusters of previously defined CA1 FMRP targets (https://hb.flatironinstitute.org/module/). Compartment-specific FMRP CLIP scores were determined essentially as described above, except without filtering for reproducibly detected mRNAs in order to maximize the number of genes included in the analysis. For GSEA analysis, CA1 mRNAs were ranked by compartment-specific FMRP CLIP scores. GSEA analysis was performed using the fgsea package (Korotkevich et al., 2021), using the gene lists from module detection as pathways.

## Acknowledgements

The authors wish to thank Alison North and Tao Tong from the Rockefeller University Bio-Imaging Resource Center for help with microscopy and image analysis, the Bioinformatic Resource Center at Rockefeller University for bioinformatics advice and support, and members of the Darnell lab for manuscript review. This work was supported by an award from the Leon Levy Foundation for Mind, Brain and Behavior to CRH, the Simons Foundation Research Award DFARI 240432 and NIH Awards NS081706 and R35NS097404 to RBD. RBD is an Investigator of the Howard Hughes Medical Institute.

---

**Supplemental Figure 1.**
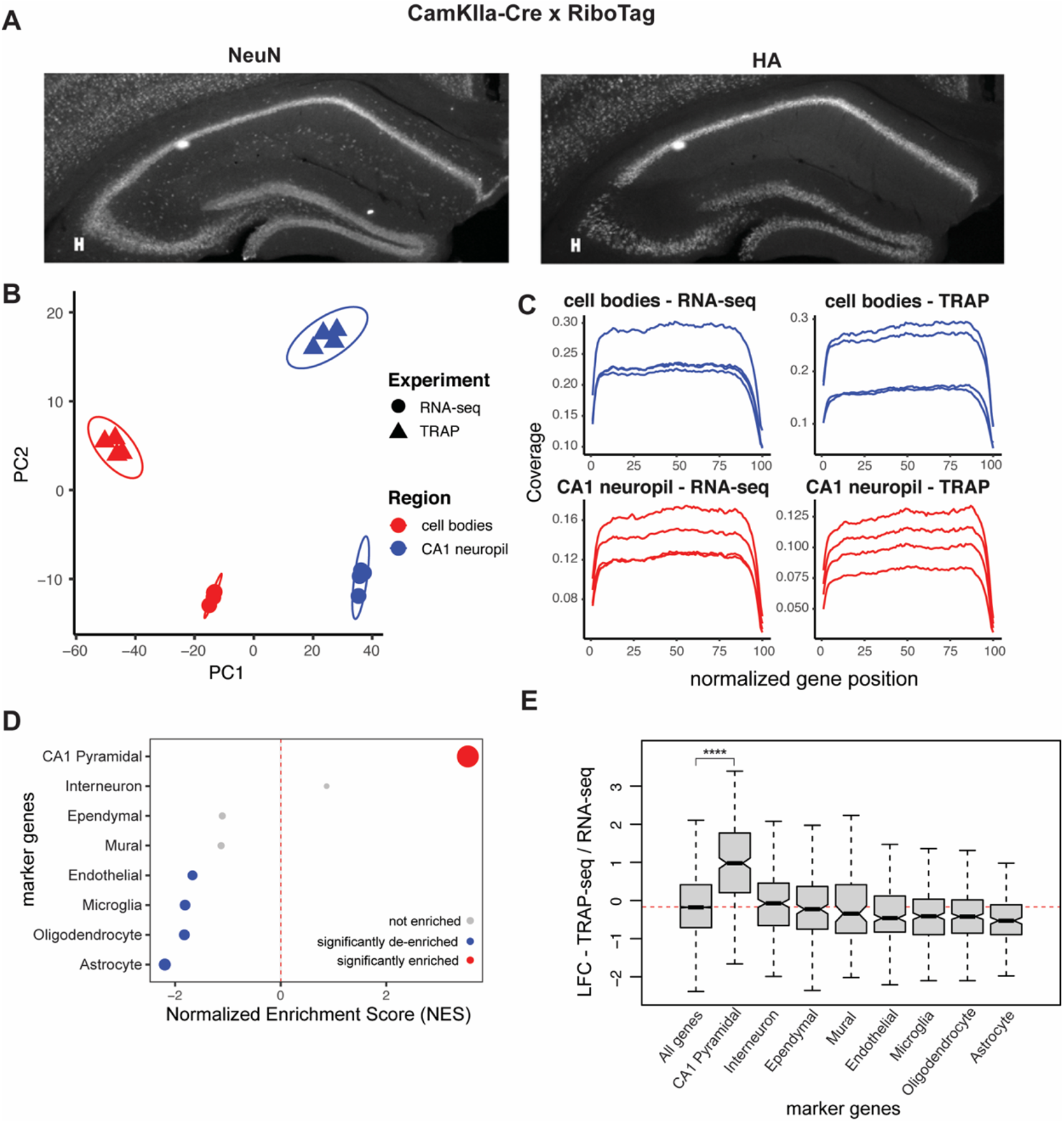
A) *Crossing the RiboTag mice with CamkIIα-Cre mice allows for expression of HA-tagged Rpl22 ribosomal subunits only in the pyramidal neurons of the hippocampus.* Immunostaining was performed on coronal brain sections from 8-10 week CamkIIα-RiboTag mice. Sections were stained with a pan-neuronal marker (NeuN, left panel) and for the HA-tagged ribosomal subunit (right panel). B) *PCA analysis of microdissected TRAP- and RNA-seq samples.* Point shape is decided by sample type (TRAP or RNA-seq) and color is determined by compartment (Cell Body Layer (CB) or CA1 neuropil) C) Gene coverage in TRAP- and bulk RNA-seq samples. Coverage of sequenced samples, as determined by Picard. D) *Contaminating cell types are not enriched in CA1 neuropil TRAP.* Genes were ranked by LFC in TRAP samples (CA1 neuropil / cell bodies) and marker gene sets were used for GSEA analysis. Significantly enriched (FDR < .05) markers are shown in red, significantly de-enriched markers are shown in blue, and not enriched cell types are shown in grey. Point size indicates -log10(FDR). E) *CA1 pyramidal markers are enriched in CA1 neuropil TRAP samples.* Boxplots compare LFC (CA1 neuropil / cell bodies) of various marker genes in CA1 neuropil TRAP. Stars indicate significance in Wilcoxon ranked sums test. (****: p-value < .00001)

**Supplemental Figure 2.**
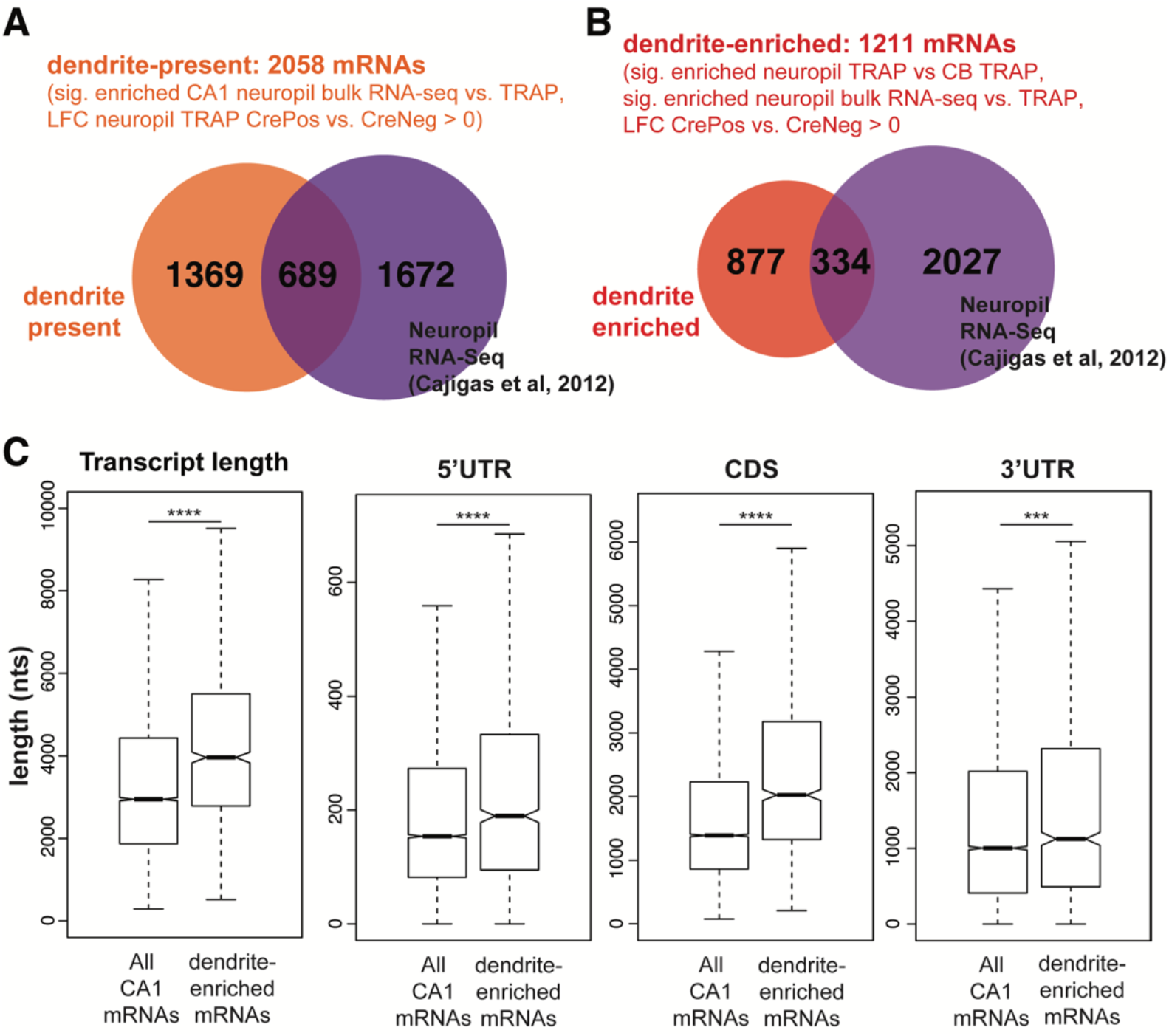
A & B) *Overlap of dendrite-present and dendrite-enriched mRNAs with previously-determined localized mRNAs by bulk RNA-seq of microdissection compartments in rat hippocampal slices* (Cajigas et al., 2012). C) *Comparison of mRNA length of all CA1-expressed mRNAs and the NE mRNAs.* The mRNA with the highest expression for each gene in CA1 neurons (determined by TRAP,) was used for this analysis. Length of dendrite-enriched and all CA1 mRNAs was compared for the full length transcript, 5’UTRs, CDS and 3’UTRs. Significance was determined by Wilcoxon ranked sums test (***: p-value < .0001, ****: p-value < .00001).

**Supplemental Figure 3.**
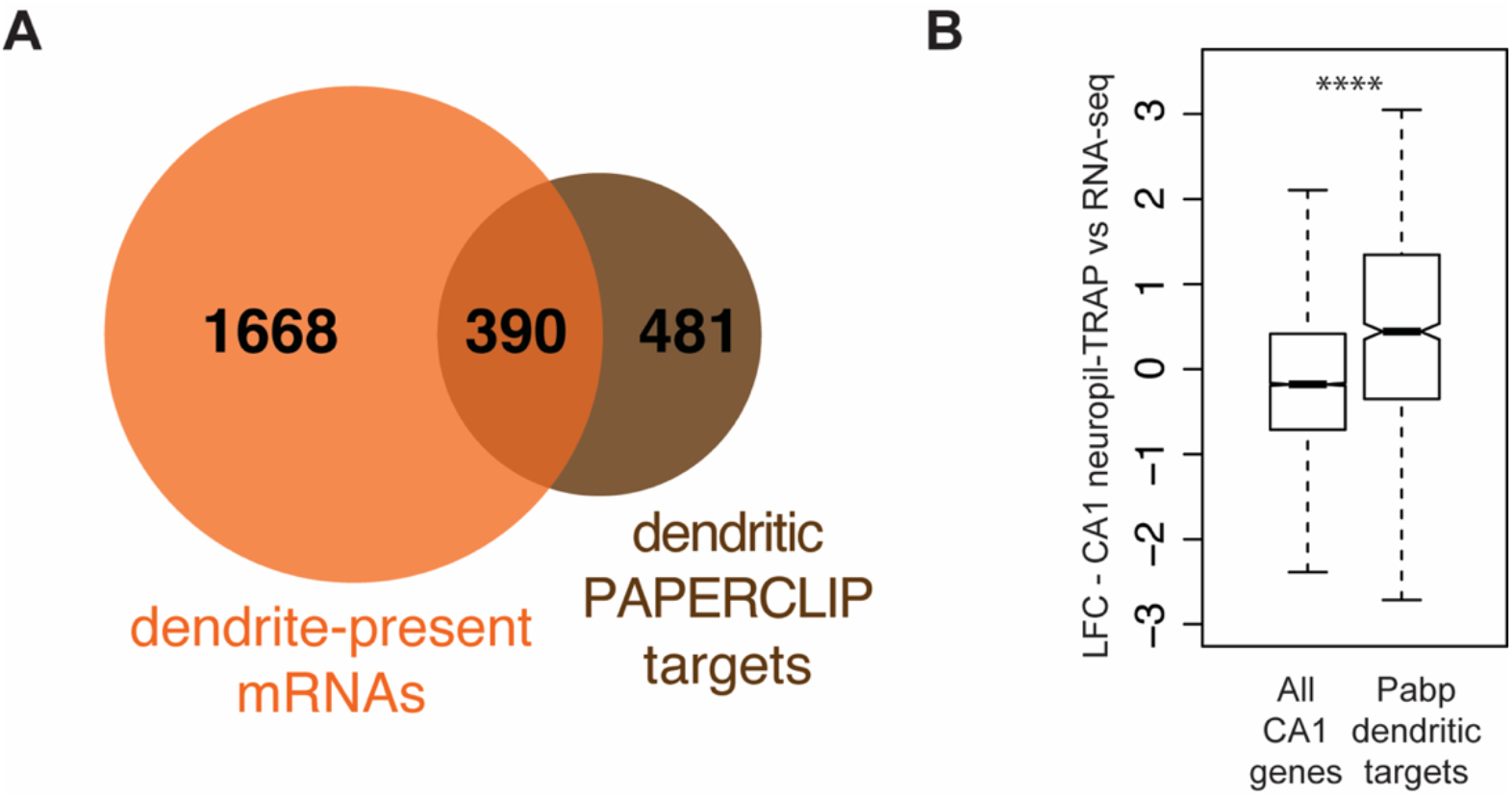
A) *Overlap of mRNAs identified by cTag-PAPERCLIP in microdissected CA1 material (orange) and dendrite-present mRNAs (brown). B) mRNAs identified by microdissected cTag-PAPERCLIP are enriched in CA1 neuropil-TRAP.* Boxplots compare the LFC (CA1 neuropil-TRAP vs bulk RNA-seq) in all CA1-expressed mRNAs and those identified by microdissected cTag-PAPERCLIP. Significance was determined by the Wilcoxon test.

**Supplemental Figure 4.**
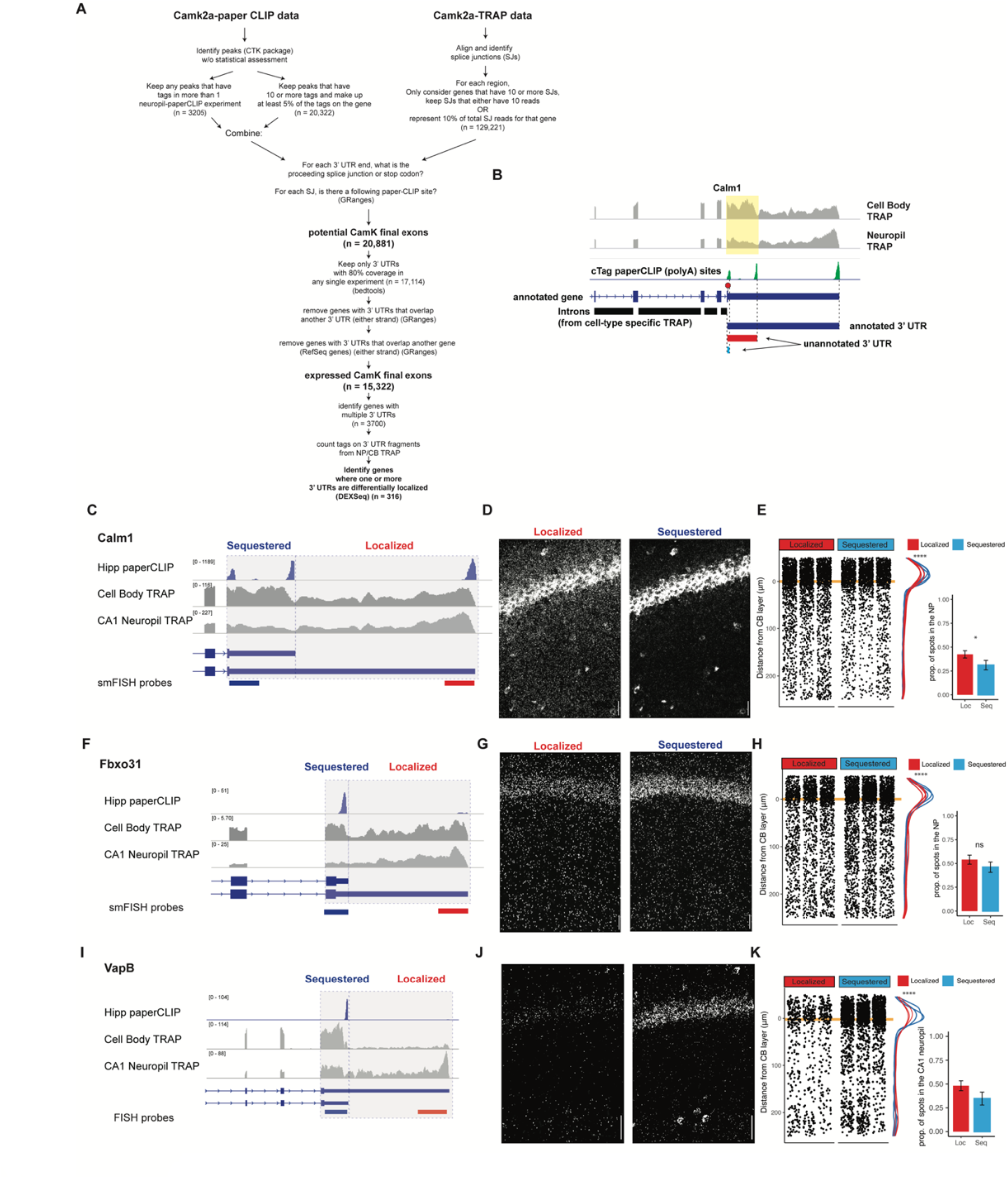
A) *Detailed schematic of 3’UTR identification pipeline. B) Identification of expressed 3’UTR boundaries by combining polyA sites from cTag-PAPERCLIP and splice junctions for cell-type specific TRAP data.* Coverage of 3’UTR regions in CA1 Cell bodies and neuropil TRAP are shown in grey. polyA sites identified by cTag-PAPERCLIP are shown in green. Introns, as determined from TRAP, are indicated by black bars. For each PAPERCLIP site (representing the 3’ end of a potential final exon), the potential 5’ boundary was determined by the nearest upstream intron (defined by TRAP). C-E) *Validation of Differential Localization of Calm1 3’UTR isoforms.* For 3’UTR-APA events (i.e. expressing a “distal” and “proximal” 3’UTR isoform), FISH probes detected “common” (found in both proximal and distal UTRs) and “specific” (found only in localized isoform) sequences. Therefore, localization of distal 3’UTR-containing mRNAs was readily detectable as decreased levels in the cell bodies of CA1 neurons. Analysis of FISH data was performed as described in Figure 2. F-H) *Validation of Differential Localization of Fbxo31 3’UTRs* I-K) *Validation of Differential Localization of VapB 3’UTRs*

**Supplemental Figure 5.**
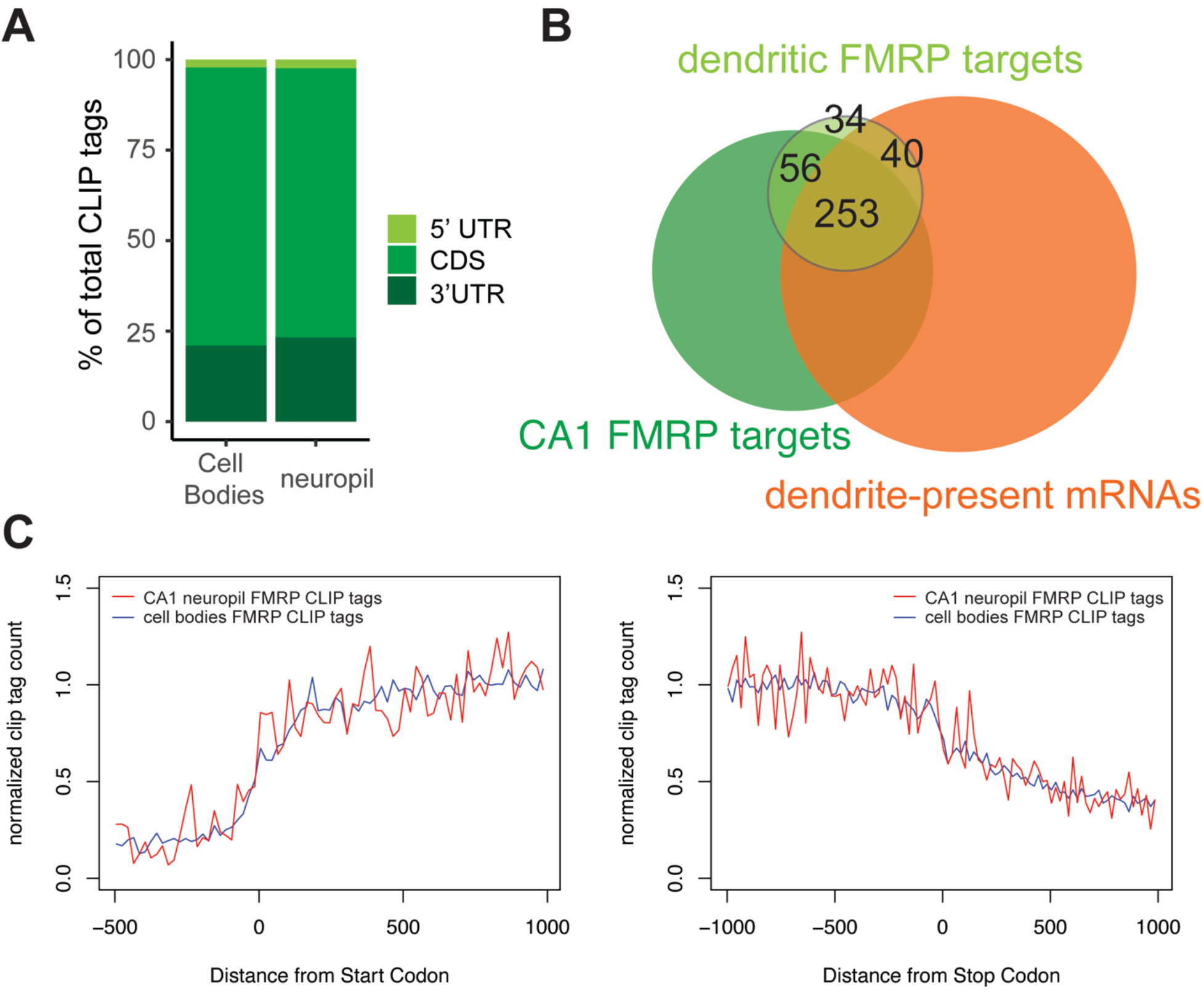
A) *Genomic distribution of FMRP cTag-CLIP tags in CA1 Cell bodies and neuropil.* Tags are aggregated for 5 biological replicates B) *Overlap between dendritic FMRP targets, CA1 FMRP targets and dendrite-present mRNAs* C) *Compartment-specific FMRP-CLIP normalized coverage across mRNAs.* Normalized tag counts are plotted for cell bodies (blue) or CA1 neuropil FMRP-cTag-CLIP (red) for the 1000 nts surrounding the start (left) or stop (right) codon.

**Supplemental Figure 6.**
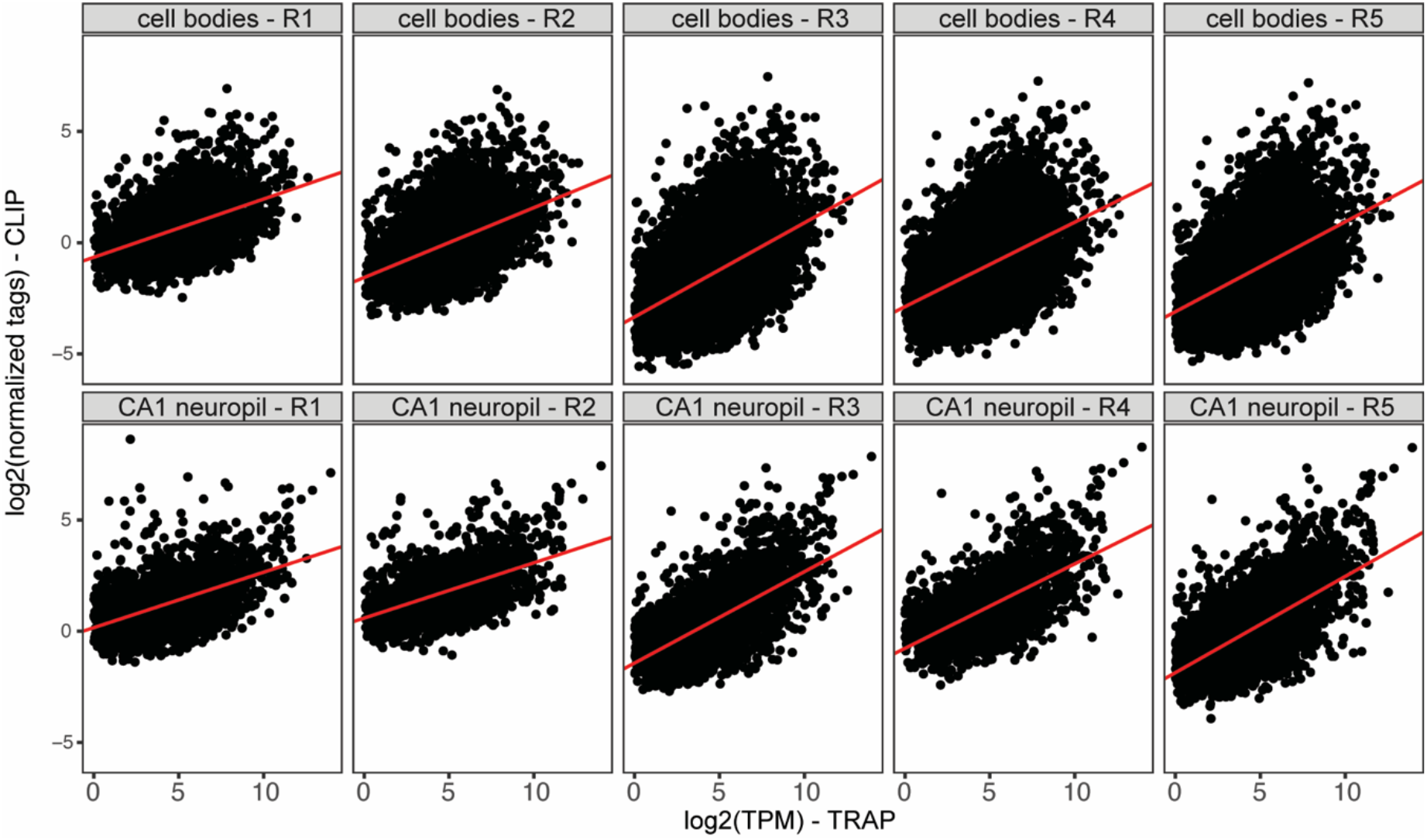
Plots represent normalized neuropil or cell bodies FMRP-CLIP tags (log2) vs log2(TPM) for cell bodies-or CA1 neuropil-CLIP and TRAP experiments. For each replicate, linear models were generated (shown in red). Compartment-specific CLIP scores are determined for each gene as the distance between each gene and the linear model line. Average CLIP scores for the five replicates were used as compartment-specific FMRP-CLIP scores for each gene.

**Supplemental Figure 7.**
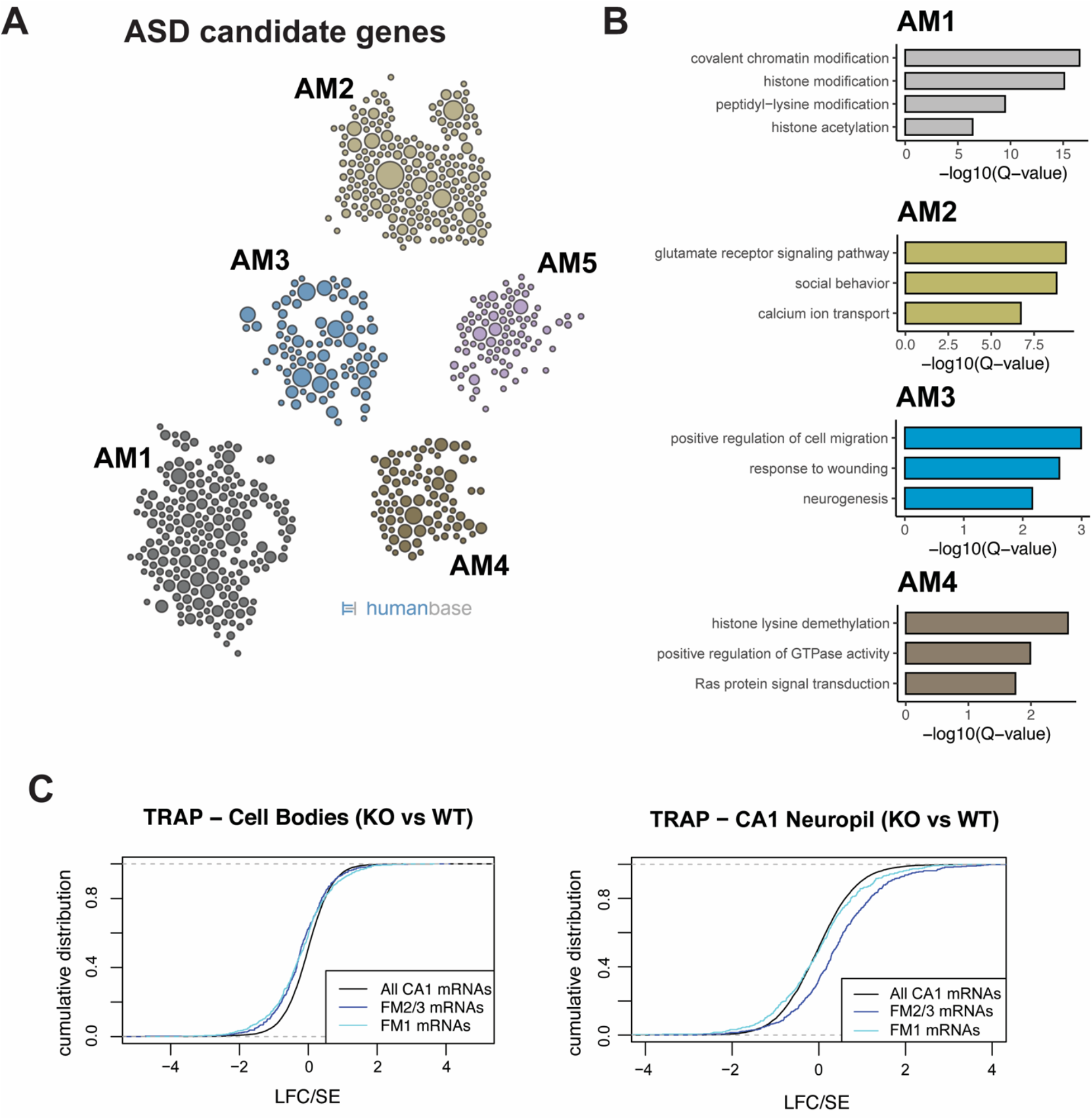
A) *Functional clusters of SFARI ASD candidate genes.* Potential ASD candidate genes as identified by SFARI were clustered with HumanBase software. B) *GO terms of top functional clusters of ASD candidate genes GO terms were identified by HumanBase software.* C) *FMRP targets are decreased in the cell bodies and increased in dendrites.* (left) FMRP target levels (LFC, KO/WT) were analyzed in cell bodies TRAP. Cdf plots are shown, and colors indicate FMRP functional clusters, as defined in Figure 7A. (right) FMRP target levels (LFC, KO/WT) were analyzed in CA1 neuropil TRAP. Cdf plots are shown, and colors indicate FMRP functional clusters.

